# The transcription factor LSL-1 interacts with the chromatin factors HIM-17, XND-1 and BRA-2 to promote the germline-specific transcriptional repertoire and to safeguard germ cell fate in *C. elegans*

**DOI:** 10.64898/2026.04.05.716469

**Authors:** Magali Nanchen, David Rodriguez Crespo, Michael Stumpe, Chantal Wicky

## Abstract

Germ cells are the only cells of an organism that pass onto the next generation and, hence perpetuate the species. To ensure this, germ cells need dedicated transcriptional repertoire, that ensure specification, proliferation, differentiation and fate maintenance. We previously characterized LSL-1, a conserved zinc-finger transcription factor that acts as a major direct transcriptional activator of genes involved in germ cell development, fate specification, meiosis and genome stability. Here, we show that LSL-1 interacts with the transcription factor HIM-17, the chromatin proteins BRA-2 and XND-1. These proteins are functionally related to LSL-1 and they colocalize at germline gene promoters, forming most likely a transcription-promoting complex. Furthermore, LSL-1 lies in close proximity to members of the COMPASS and the MOF complexes, corroborating the observation that HIM-17 and LSL-1 are required to maintain normal level of H3K4 methylation in the gonad. Finally, we show that LSL-1 interacting partners are necessary to maintain germ cell fate. Altogether, we propose that LSL-1 interacts with transcription regulators and chromatin modifiers to ensure the establishment of the transcriptional repertoire appropriate for germline function as well as for cell fate maintenance.

## Introduction

The germline is a special population of cells that is essential for reproduction and perpetuation of the species. They required dedicated gene regulation at transcriptional and post-transcriptional levels to get specified, to proliferate and to differentiate into gametes (Strome and Updike, 2015; Wang and Seydoux, 2013). Finally, germ cell fate needs protective mechanisms to be maintained and to allow birth of the next generation (Strome and Updike, 2015). Germline gene repertoires have been characterized in several organisms, including the nematode *C. elegans*, and they include transcriptional regulators, chromatin factors, meiotic genes and post-transcriptional regulators (Fierro-Constaín et al., 2017; Reinke et al., 2004). In *C. elegans* germline gene regulation occurs mostly regulated at the post-transcriptional level (Merritt et al., 2008). However, we identified recently the transcription factor LSL-1 that is required to produce functional gametes. *lsl-1* encodes a zinc-finger transcription factor, that is specifically expressed in the germ cells throughout development, from the P4 blastomere to adult germ cells. *lsl-1* encodes a protein with three well conserved N-terminal C2H2-type zinc-finger domains (Rodriguez-Crespo et al., 2022). The three N-terminal zinc fingers closely resemble to those that characterize the SP1/Krüppel-like family of transcription factors, a protein family with diverse functions in growth and development (Kaczynski et al., 2003; Pearson, 2013). Transcriptomic and chromatin binding analysis indicate that LSL-1 activates the germline transcriptional repertoire by binding to the promoter of genes involved in germ cell proliferation, meiotic pairing and recombination, genome stability and P granule composition (Kudron et al., 2018; Rodriguez-Crespo et al., 2022). Identification of LSL-1 as a major regulator of germline gene expression raises the question of its role in germ cell fate maintenance.

Extensive studies on the nematode *C. elegans* have allowed molecular characterization of germ cell fate protective mechanisms, that are conserved in other species. Germ cell immortality is ensured by barriers to somatic reprogramming that range from translational control to chromatin regulation and efficient DNA repair mechanisms (reviewed in Fassnacht and Ciosk, 2017; O’Neil and Rose, 2006; Robert et al., 2015; Ul Fatima and Tursun, 2020). Particularly relevant to this study is the role of chromatin as a barrier to prevent reprogramming. Altering H3K4 level of methylation leads to spontaneous reprogramming of germ cells. Our study shows that lack of SPR-5/LSD1 H3K4 demethylase activity leads to teratoma formation in the germline (Käser-Pébernard et al., 2014). Intriguingly, perturbation of the opposite activity, namely the H3K4 methyltransferase SET-2 and other subunits of the COMPASS complex responsible for global H3K4 methylation, also leads to expression of somatic genes in the germline and to transdifferentiation of germ cells (Robert et al., 2014). In this study, reprogramming of germ cells is accompanied by a global reorganization of the chromatin, including a decreased level of the repressive mark H3K9 methylation. Altogether these results suggest a complex, locus specific, chromatin organization that deserve to be further studied.

Here, we show that LSL-1 interact with transcription factors and chromatin proteins to activate germline genes and to ensure germ cell fate maintenance. Mass-spectrometry analysis of the LSL-1 complex and neighboring proteins residing revealed that LSL-1 forms a core complex with the THAP-domain protein HIM-17, the XND-1 chromatin protein and the MYND domain protein BRA-2. It also lies in proximity to members of the COMPASS and the MOF complexes, both modifying chromatin in a way to favor transcription. Furthermore, we show that LSL-1, as well as core interactors prevent germ cells to be converted into neuron-like cells. We propose that LSL-1 and interactors are activating germline gene transcription by targeting the COMPASS and the MOF complex at germline gene promoters. This leads to proper H3K4me3 deposition and to germ cell fate maintenance.

## Results

### LSL-1 interacts with transcription regulators

To better understand how LSL-1 is acting on transcription, we determined its protein interaction network by performing coimmunoprecipitation followed by mass spectrometry analysis (CoIP-MS). IP was performed using a strain expressing *lsl-1::gfp* and a GFP-trap bead system. To extend the study to protein residing in the vicinity of LSL-1, we also determined the protein proximity network of LSL-1. Proximity labeling was achieved by fusing LSL-1 to the TurboID variant of the biotin ligase (Artan et al., 2021; Branon et al., 2018). The expression pattern of the LSL-1::TurboID protein corresponds to the LSL-1 protein (Figure S1) (Rodriguez-Crespo et al., 2022). Volcano plots show the intensity/abundance of LSL-1 and its interacting partner or nearby protein peptides compared to the control samples – GFP-control strain for CoIP-MS and wildtype control strain for TurboID-MS – on the x-axis (Figure 1A and 1 A’, Sup Table 1 and 2). Among the significant hits, we focused on candidates represented by more than one peptide, leading to 203 LSL-1 proximal proteins and 194 LSL-1 interactors (Figure 1B). Among them, 16 were found in common (Figure 1B), including three of the most enriched proteins in the interactome, namely the MYND-domain protein BRA-2, the chromatin protein XND-1 and DPY-30, which is involved in dosage compensation. The interaction between LSL-1 and XND-1 was validated by co-immunoprecipitation and XND-1 detection on western blot (Figure S1).

**Figure 1.**
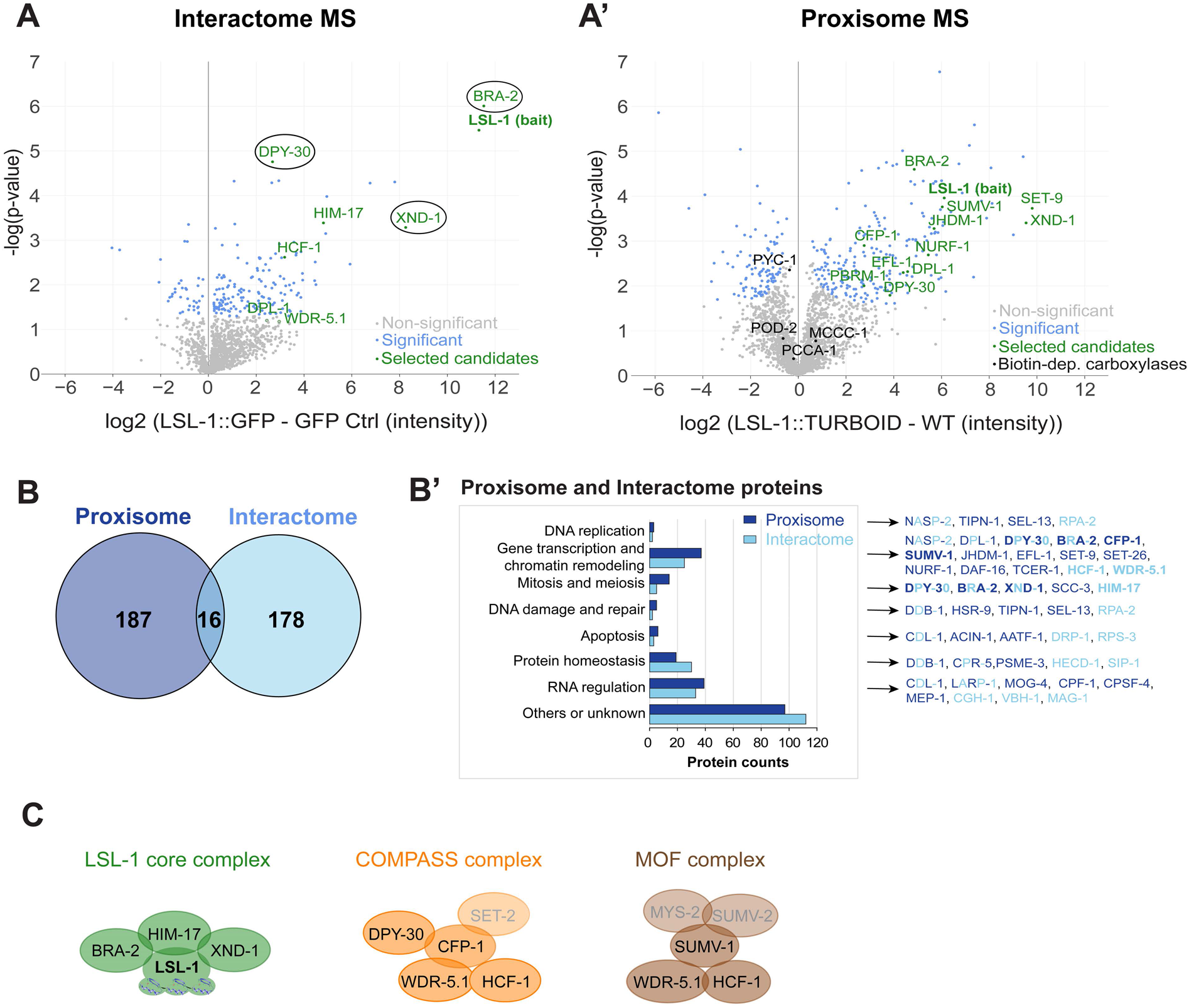
LSL-1 protein interaction network. (A) Volcano plots show the log2 fold change of protein intensities in the LSL-1-tagged sample compared with the control sample (x-axis) versus the –log10 p-value (y-axis). CoIP-MS analysis is on the left and the proximity-labeling-MS analysis on the right. The threshold was set at p-value < 0.05. (A, A’) Grey are non-significant hits, blue and green are significant hits. (A’) Black are the endogenous biotin-dependent carboxylases. Black circles highlight the three most-enriched proteins in the interactome, also found in the proxisome. (B) Common enriched proteins which p-value < 0.05 and which are represented by at least more than one peptide. Dark blue, proxisome. Light blue, interactome. (B’) Functional annotation of enriched proteins found in CoIP-MS and TurboID-MS data sets according to wormbase, with examples for each category on the left. Dark blue, proxisome. Light blue, interactome. Bold is used for the proteins in the selected complexes. (C) Protein complexes interacting with LSL-1. Protein names in grey belong to complex members not found in the MS. Green, LSL-1 core complex. Orange, COMPASS complex. Braun, MOF complex.

Using the SimpleMine tool from Wormbase, functions associated with gene transcription, chromatin remodeling, protein homeostasis, and RNA regulation were over-represented among the interactors (Figure 1 B’). HIM-17, is known to activate transcription of germline genes and BRA-2 as well as XND-1 are chromatin proteins active in the germline, in particular during meiotic prophase. (Bessler et al., 2007; Blazickova et al., 2025; Nadarajan et al., 2021; Reddy and Villeneuve, 2004; Wagner et al., 2010). Based on interaction intensities and functional data, we propose that LSL-1 forms a core protein complex with BRA-2, HIM-17, and XND-1 chromatin.

Interestingly, we identified, mainly in the proxisome but also in the interactome, members of the COMPASS complex (WDR-5.1, HCF-1, DPY-30 and CFP-1) responsible for H3K4 methylation and the MOF histone acetyltransferase complex (WDR-5.1, HCF-1 and SUMV-1) Both complexes are known to be involved in chromatin remodeling and transcription activation (Figure 1C) (Herbette et al., 2020; Hoe and Nicholas, 2014; Li and Kelly, 2011). Altogether, this indicates that the LSL-1 core complex might interact with the COMPASS and the MOF to mediate transcriptional activation.

Additional proximal proteins related to chromatin remodeling were uncovered. SET-9 is an H3K4me3 reader that restricts the spreading of this histone mark and is necessary for proper germline development (Ni et al., 2012; Wang et al., 2018). JHDM-1 is a putative H3K9me2 demethylase known to play a role in aging (Lee et al., 2019). NURF-1, a member of the NURF complex, promotes the opening of the chromatin through the binding to H3K4me3 (Large et al., 2016). All these proteins contribute to chromatin dynamics through H3K4 and H3K9 methylations and might function with LSL-1 to activate transcription. Finally, the transcription factor DPL-1 and its known partner EFL-1 were also identified as LSL-1 interactor and proximal protein, respectively. Their role in promoting oogenesis and embryogenesis, by inducing the transcription of tissue-specific genetic programs is consistent with an interaction with LSL-1 (Ceol and Horvitz, 2001; Chi and Reinke, 2009, 2006).

### Core complex members are binding to common chromatin regions via LSL-1

To determine whether LSL-1 and its core interacting partners XND-1 and HIM-17 are binding to chromatin as a complex, we analyzed and compared ChIP-seq data from the Modern Consortium and from the Ahringer lab, obtained from young adult worms expressing either LSL-1::TY1::EGFP::3xFLAG (*wgls720*) or XND-1::EGFP::3xFLAG transgenes (Carelli et al., 2022; Kudron et al., 2018). HIM-17 ChIP-seq data were obtained using an anti-HIM-17 antibody. We looked at XND-1 and HIM-17 enrichment at the 3’896 identified LSL-1 binding sites (2kb intervals) (Kharchenko et al., 2008) (Figure 2A). The resulting heatmaps highlight a strong enrichment of HIM-17 and XND-1 at LSL-1 binding sites, supporting the idea that LSL-1, XND-1, and HIM-17 are forming a complex functioning at shared chromatin regions.

**Figure 2.**
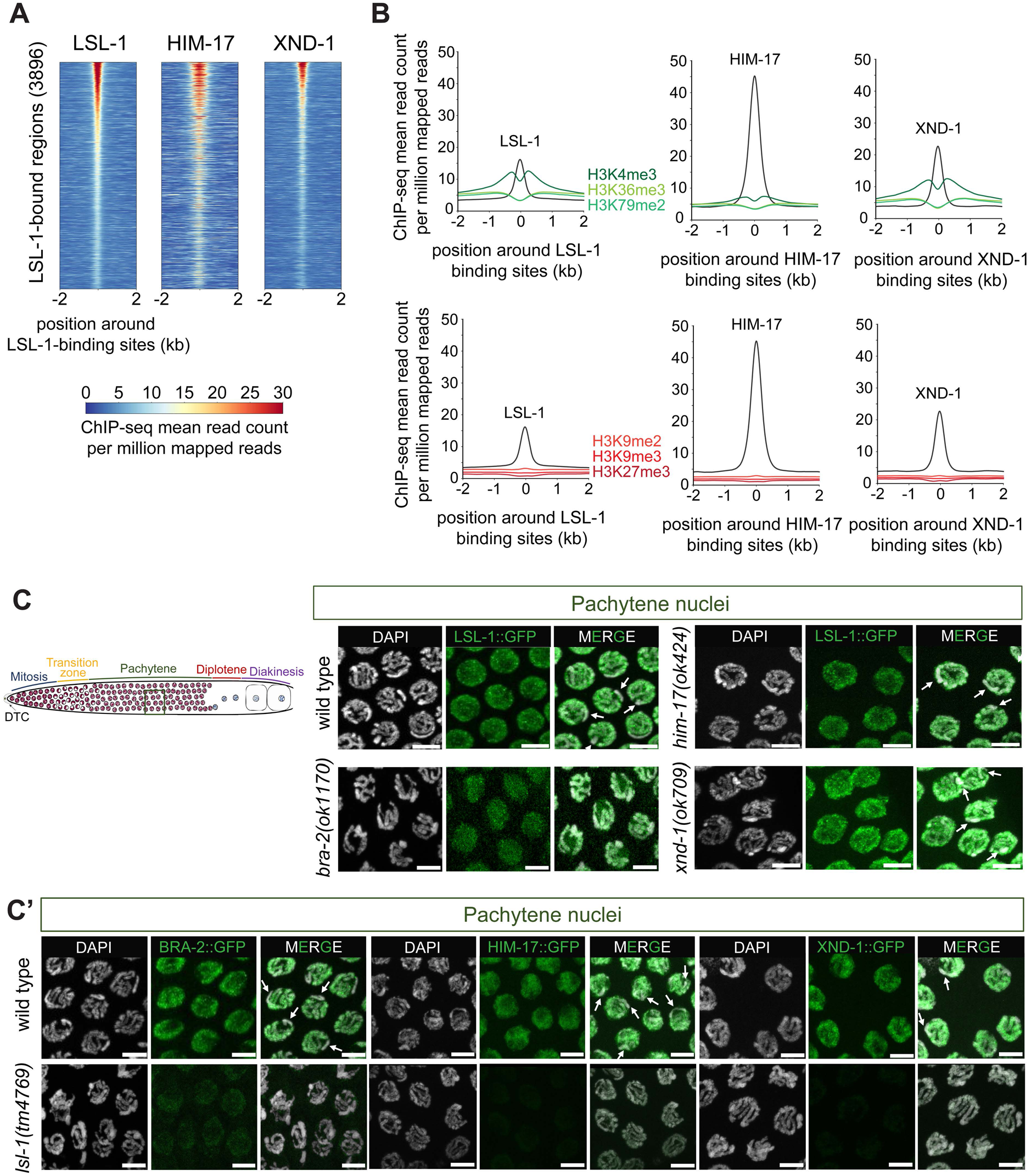
LSL-1, HIM-17, and XND-1 bind to shared chromatin regions that are associated with active chromatin marks. (A) Heatmaps show enrichment of LSL-1, HIM-17, and XND-1 at the LSL-1-bound regions determined by ChIP-seq peaks (3896). (B) Profile plot shows the LSL-1, HIM-17, or XND-1 binding regions enrichment of LSL-1, HIM-17, XND-1, H3K4me3, H3K36me3, H3K79me2, H3K9me2, H3K9me3, and H3K27me3, centered on LSL-1, HIM-17, or XND-1 binding regions (ChIP-seq peaks). (A, B) The reads were positioned around the center of the LSL-1 binding sites with 2 kb regions upstream and downstream. Representative confocal projection images of pachytene nuclei of 1-day-old adult stage gonads of (C) wild type, *bra-2(ok1170)*, *him-17(ok424)* or *xnd-1(ok709)* mutans, expressing LSL-1::GFP and stained with DAPI and of (C’) wild type and *lsl-1(tm4769)* mutants, expressing BRA-2::GFP, HIM-17::GFP, or XND-1::GFP and stained with DAPI. Scale bars, 5 µm.

Using ChIP-seq data from the Modern consortium, we looked at DPL-1 and EFL-1 enrichment at LSL-1 binding regions (Kudron et al., 2018). While DPL-1 is only mildly enriched, EFL-1 shows a strong enrichment, indicating that it might also function with LSL-1 in a subset of chromatin regions (Figure S2A). By selecting regions where LSL-1 binds in the promoters, HIM-17, XND-1 and DPL-1 show even a higher enrichment at LSL-1 binding sites, indicating that these proteins are functioning together in promoters (Figure S2B).

To further analyze the chromatin context at LSL-1, HIM-17, and XND-1 binding sites and the link with the COMPASS and/or MOF complexes, we used available ChIP-seq data for active chromatin marks (Jänes et al., 2018; McManus et al., 2021, modEncode_5169 project for H3K79me2) and for repressive marks (McMurchy et al., 2017; Zaghet et al., 2021). LSL-1 binding regions show a strong enrichment in H3K4me3 and the same trend was observed for XND-1. However, H3K36me3 and H3K79me2 were only mildly enriched at LSL-1 and XND-1 binding sites. HIM-17 binding regions present almost no association with the histone modifications analyzed here. In contrast, repressive marks such as H3K9me2, H3K9me3, and H3K27 are largely absent from the binding regions of all three proteins (Figure 2 B).

Taken together, these results support the idea that LSL-1 functions as a transcriptional activator, together with HIM-17 and XND-1. It may promote gene activation by interacting with the COMPASS chromatin-remodeling complex.

To investigate how the LSL-1 core complex members rely on each others for recruitment at the chromatin, we monitored their localisation in various mutant backgrounds. In wild-type worms, LSL-1, BRA-2, HIM-17, and XND-1 share a similar localization pattern. They are associated to the autosomes and to the nucleoplasm of germ cells (Figure 2C and C’) (Blazickova et al., 2025; Reddy and Villeneuve, 2004; Rodriguez-Crespo et al., 2022; Wagner et al., 2010).

In the absence of BRA-2, the LSL-1::GFP signal appears more diffused, indicating that LSL-1 association with the chromatin is perturbed (Figure 2 C). BRA-2 might be required for the stabilization of LSL-1 binding at the chromatin. In contrast, LSL-1 localization is not altered by the absence of HIM-17 or XND-1, indicating that these two proteins are not necessary for LSL-1 recruitment at the chromatin (Figure 2 C).

BRA-2 is detected in germline as well as in somatic nuclei, including neuronal nuclei (Figure 2 C’ or Figure S2). In the absence of LSL-1, BRA-2 is barely detectable in germline nuclei but remains present in somatic nuclei. HIM-17 and XND-1 also disappear from the chromatin when LSL-1 is missing, indicating that LSL-1 is required for the localization of BRA-2, HIM-17 and XND-1 at the germline chromatin (Figure 2 C’).

Altogether, these results indicate that LSL-1 is the main recruiter of the complex.

### LSL-1 and HIM-17 are regulating a large set of common direct target genes

To determine whether LSL-1, HIM-17, and XND-1 could function as a complex in transcription regulation, we analysed and compared available ChIP-seq and transcriptomic data of the core complex members (Carelli et al., 2022; Kudron et al., 2018; Rodriguez-Crespo et al., 2022). LSL-1 and HIM-17 bind to a similar number of genes: 3’552 for HIM-17 and 3’273 for LSL-1, and 1’758 are common, of which 77.7% show binding in the promoter region (Figures 3 A, Figure S3). XND-1 binds to 1’646 genes, most of them being bound by LSL-1 (86.6%). 81.2% of them are occupied by LSL-1 and XND-1 in the promoter regions (Figure 3A’). Overlapping genes that are bound by the three factors uncovered 997 genes (Figure 3 B). These results indicate that LSL-1, HIM-17 and XND-1 are binding to a large set of genes (Figure 3 B).

**Figure 3.**
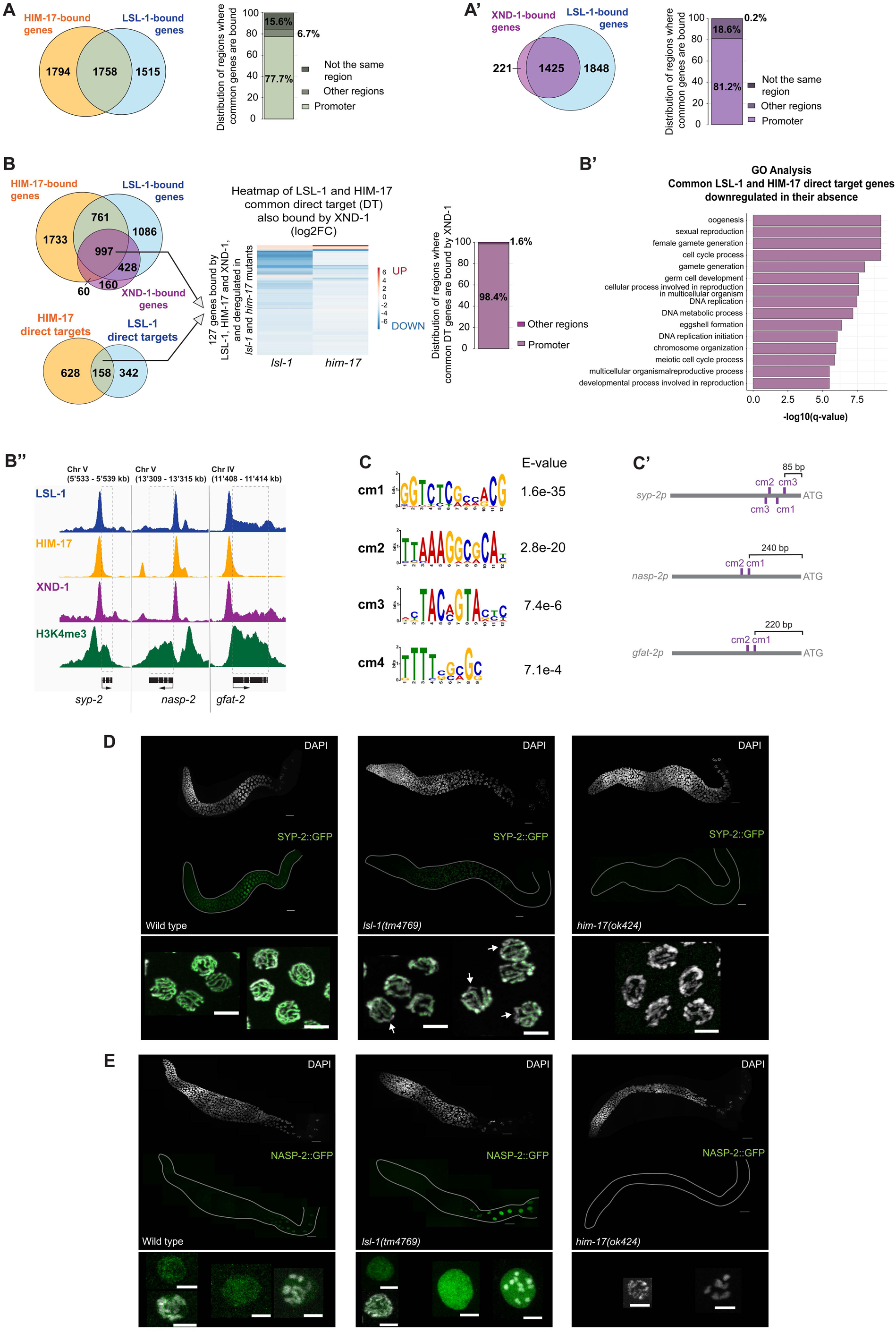
LSL-1, HIM-17, and XND-1 bind to shared target genes, part of which are activated by LSL-1 and HIM-17. (A and A’) Overlap between LSL-1 (blue) and HIM-17 (orange) or XND-1 (purple) bound genes, with stacked plots showing the distribution of genomic regions bound by LSL-1 and HIM-17 or XND-1. (B) Overlap between LSL-1 (blue), HIM-17 (orange) and XND-1 (purple) bound genes and overlap between *him-17* (orange) and *lsl-1* (blue) DEGs targeted by both HIM-17 and LSL-1. Heatmap illustrates the log2Foldchange of 127 DEGs common to *lsl-1* and *him-17* mutants and also bound by HIM-17, LSL-1 and XND-1, red indicates upregulated genes, and blue downregulated ones. The stacked plot shows the distribution of genomic regions bound by XND-1 for these 127 direct target genes. (B’) Bar plots represent the Gene Ontology enrichment analysis of the common gene lists. (B’’) IGV genome viewer screenshots (Robinson et al., 2011; Thorvaldsdóttir et al., 2013) depicting enrichment of LSL-1, HIM-17, XND-1 and H3K4me3 at *syp-2*, *nasp-2* and *gfat-2* promoters. (C) Sequence logo of significant motifs enriched at the promoter of both HIM-17 and LSL-1 direct target genes, identified using the MEME-platform (MEME version 5.5.9). (C’) Schematic drawing of promoter regions of *syp-2*, *nasp-2* and *gfat-2* genes including the common MEME binding motifs (cm). (D and E) Representative confocal projection images of 1-day-old adult gonads of wild type, *lsl-1(tm4769)* and *him-17(ok424)* worms, expressing SYP-2::GFP (D) or NASP-2::GFP (E) and stained with DAPI. Gonad shapes were delimited by contouring for the GFP channel. Lower panel: magnification of pachytene and diakinesis germline nuclei. Scale bars, 20 µm for whole gonad representation and 5 µm for each magnification.

Based on transcriptomic data, we identified 895 genes that were differentially expressed in both, *him-17* and *lsl-1* mutants, 93.6% of them being downregulated (Carelli et al., 2022; Rodriguez-Crespo et al., 2022) (Figure S3). Overlapping the ChIP-seq and transcriptomics data delivered 127 genes that are directly regulated by LSL-1, HIM-17 and XND-1, out of which 91.8% are being downregulated in absence of HIM-17 and LSL-1 activity (Figure 3 B, Sup Table 3). Go term analysis of this gene set suggest that LSL-1, HIM-1 and XND-1 function together as a complex at promoter of genes involved in reproduction and genome organization (Figure 3 B-B’). By inspecting the promoter regions of three direct target genes, namely *syp-2*, *nasp-2* and *gfat-2*, we observed that LSL-1, HIM-17 and XND-1 were bound in the same region, also enriched in H3K4me3 (Figure 3 B’’). Using the MEME tool, we could identified in LSL-1, HIM-17 and XND-1 target genes, including *syp-2*, *nasp-2* and *gfat-2*, the motifs that were previously proposed to be recognized by HIM-17 and LSL-1 (Bailey et al., 2015; Bailey and Elkan, 1994; Carelli et al., 2022; Rodriguez-Crespo et al., 2022) (Figure 3 C-C’). These results support the idea that LSL-1, HIM-17 and XND-1 are binding in form of a complex at promoter regions enriched with active chromatin marks.

To measure the impact of LSL-1 and HIM-17 on transcription *in vivo*, we generated transgenic animals expressing *nasp-2* and *syp-2* tagged with GFP (Figure 3 D-E). s*yp-2* and *nasp-2* encode a transversal element of the synaptonemal complex and a histone chaperone, respectively (Colaiácovo et al., 2003; Cook et al., 2011; Horard and Loppin, 2015; Smolikov et al., 2007; Tirgar et al., 2022). We analyzed the GFP localization in wild-type, *lsl-1(tm4769)* and *him-17(ok424)* mutants. In wild-type worms, SYP-2::GFP localisation corresponds to the one previously characterized (Colaiácovo et al., 2003; Schild-Prüfert et al., 2011). SYP-2::GFP appears in transition zone remains associated with the chromosomes until the end of meiotic prophase (Figure 3 D). In *lsl-1(tm4769)* mutants, overall GFP intensity is decreased, and some regions of the chromatin are depleted of SYP-2. In *him-17* mutants, we could not detect any GFP signal (Figure 3 D). These results confirm that *syp-2* is regulated by both LSL-1 and HIM-17. In wild-type worms, NASP-2::GFP appears in diplotene nuclei and remains present in oocyte nuclei. NASP-2::GPP localizes to the chromatin as well as to the nucleoplasm (Figure 3 E). Surprisingly, NASP-2::GFP is not depleted from *lsl-1(tm4769)* mutant (Figure 3 E), while it is totally absent from *him-17(ok424)* mutant gonads (Figure 3 E). In contrast to the global *nasp-2* mRNA level, which is decreased in *lsl-1(tm4769)* (-2.29 log2Foldchange), the NASP-2::GFP protein level remains the same in diplotene nuclei and even appears increased in oocyte nuclei compared to wild-type (Rodriguez-Crespo et al., 2022). One simple interpretation is that despite a lower global mRNA level in the *lsl-1(tm4769)* mutant, post-translational regulation ensures a protein amount comparable to the wild-type level or even higher in oocyte nuclei. Alternatively, expression of *nasp-2* from the transgene does not recapitulate the endogenous pattern of expression and might lack some regulatory elements. However, LSL-1 and HIM-17 binding sites are present on the transgene.

### LSL-1 interactors and neighbors are functionally related

Next, we analysed whether LSL-1 interactors and neighbors are functioning in the same biological processes. LSL-1 and the core complex members, BRA-1, XND-1 and HIM-17, are localized in germline nuclei (Blazickova et al., 2025; Reddy and Villeneuve, 2004; Rodriguez-Crespo et al., 2022; Wagner et al., 2010 and Figure 2 C and C’). BRA-2 is also detected in somatic nuclei (Figure S2). Genes encoding members of the COMPASS and the MOF complexes, including *dpy-30*, *cfp-1*, *wdr-5.1*, *hcf-1* and *sumv-1*, are expressed in germline and somatic tissues (Beurton et al., 2019; Hsu et al., 1995; Kumari et al., 2023; Li et al., 2008; Li and Kelly, 2014; Yücel et al., 2014; Zeng et al., 2026) (Figure 4 A).

**Figure 4.**
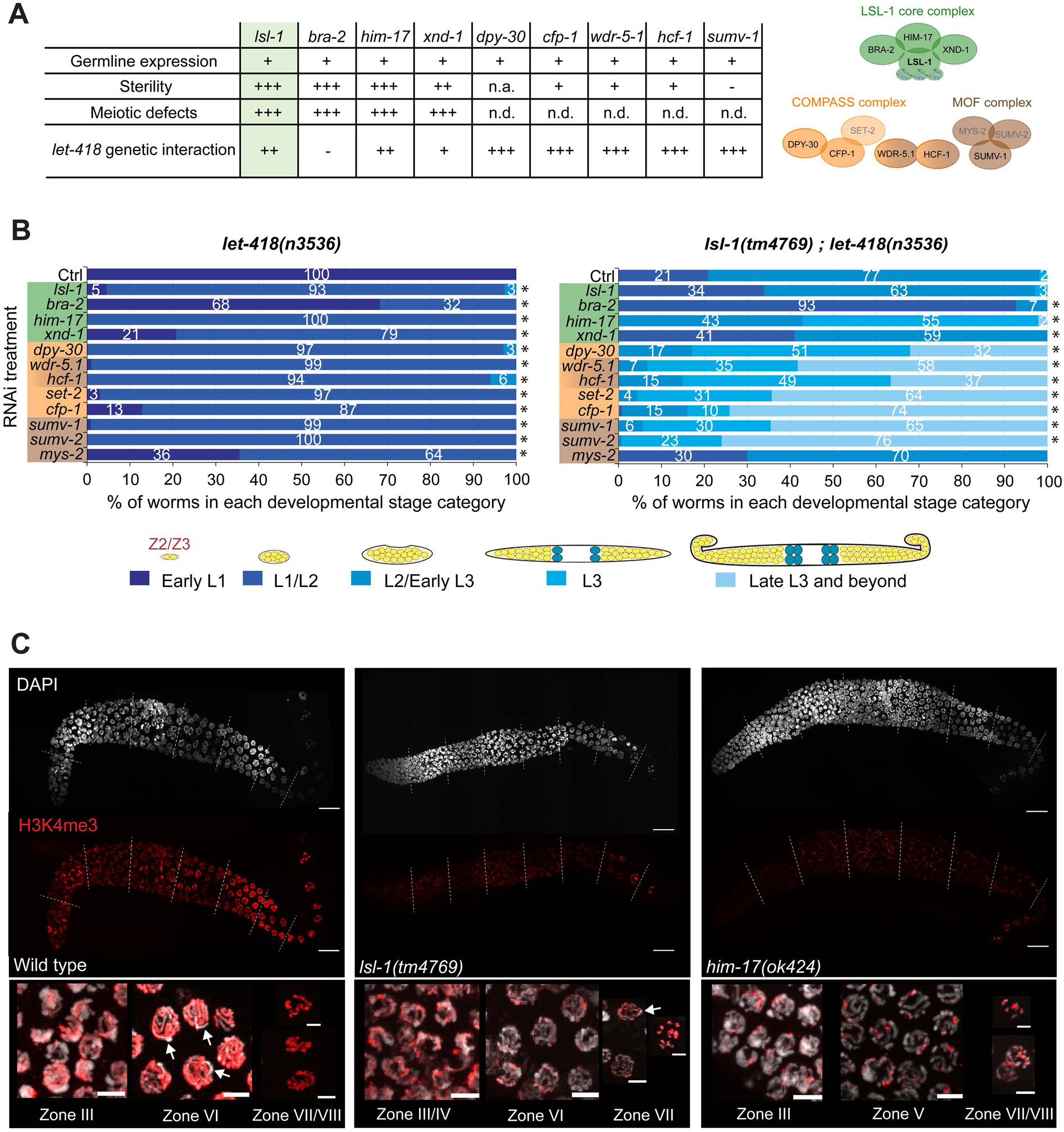
Functional relationship between LSL-1 interactor and neighbors. (A) Summary table of the features associated with *lsl-1* and its interactors. The number of + indicates presence and intensity of the feature. n.a. stands for not applicable, while n.d. for not determined. (B and B’) Stacked bar plot showing the percentage of *let-418(n3536)* or *lsl-1(tm4769);let-418(n3536)* progeny at different developmental stages. The color-coded drawings of the gonad developmental stages are depicted below the bar plot (see also Figure S4). LSL-1 core, COMPASS and MOF complex members are highlighted in green, orange, and brown, respectively. Statistical tests were performed through a Fisher–Freeman–Halton test between each RNAi condition and the control strain. Stars indicate a p-value < 0.05. (C) Representative confocal projection images of 1-day-old adult gonads of wild type, *lsl-1(tm4769)* and *him-17(ok424)* worms, stained with H3K4me3 antibodies and DAPI. Dashed lines delineate the eight equivalent-sized zones of the gonad. Below is a magnification of the germline nuclei in the indicated zones. Arrows point at the X chromosome. Scale bars, 20 µm for whole gonad representation and 5 µm for each magnification.

Mutations in genes encoding core complex members lead to sterility associated with defects in meiotic prophase, including meiotic recombination and chromosome pairing. (Bessler et al., 2007; Blazickova et al., 2025; Mainpal et al., 2015; McClendon et al., 2016; Nadarajan et al., 2021; Raices et al., 2025; Reddy and Villeneuve, 2004; Rodriguez-Crespo et al., 2022; Wagner et al., 2010). Members of the COMPASS or the MOF complexes CFP-1, WDR-5.1 and HCF-1 are also required for worm fertility, although their function in meiotic processes was not demonstrated (Lee et al., 2007; Li and Kelly, 2011; Pokhrel et al., 2019; Zeng et al., 2026). Mutations in *dpy-30* are lethal and prevent any phenotypic analysis of the gonad (Hsu et al., 1995; Hsu and Meyer, 1994). Altogether, gene expression data and corresponding mutant phenotype analysis agree with LSL-1 interactors playing a role in common biological processes.

To further investigate LSL-1 interactor functions, we tested their ability to antagonize LET-418 function. Lack of *lsl-1* activity was shown to suppress ectopic expression of germline genes in the soma, as well as developmental defects associated with mutations in *let-418* (Erdelyi et al., 2017). Thus, to assay this genetic interaction with LSL-1 interactors, we treated *let-418* mutants, as well as *lsl-1;let-418* double mutants, expressing the germ cell specific PGL-1::mCherry transgene, with RNAi corresponding to core interactors, COMPASS and MOF members (Figure 4 B and Figure S4). To get a better insight in to the interaction of *lsl-1* with members of the COMPASS and the MOF complexes, we included in the RNAi treatment *set-2*, *sumv-2* and *mys-2*. We uncovered that RNAi treatment corresponding to all candidate interactors significantly suppress, however at different levels, the L1 developmental arrest of *let-418(n3536)* mutant, indicating that they function genetically as *lsl-1* (Figure 4 B). Interestingly, *set-2*, *sumv-2* and *mys-2* RNAi treatments were also suppressing *let-418* developmental arrest.

Next, we applied the same RNAi treatments to the *lsl-1(tm4769);let-418(n3536)* double mutants. *bra-2(RNAi)* worsen the phenotype, suggesting that *bra-2* might also function independently of *lsl-1* to promote post-embryonic development. *xnd-1(RNAi)* and *mys-2(RNAi)* do not enhance the development of *lsl-1(tm4769);let-418(n3536)* mutant larvae, indicating that LSL-1 are functioning together with XND-1 and MYS-2. Finally, *him-17(RNAi)*, *sumv-1(RNAi)*, *sumv-2(RNAi)* and COMPASS member RNAi treatment, trigger a synthetic effect with *lsl-1* in suppressing the developmental arrest of *let-418* mutants, with a subset of worms reaching adulthood. This indicates either that LSL-1 and these proteins are functioning in parallel pathways to antagonize LET-418 function or that removing more than one of the proteins is more deleterious to the function of the complexes. Overall, RNAi of the core complex members has a lower impact on the development of *lsl-1;let-418* mutants compared COMPASS and MOF components, reinforcing the idea that BRA-2, XND-1 and HIM-17 are functioning as a core complex.

Next, we examined the role of LSL-1 and HIM-17 in H3K4me3 marks deposition or maintenance. To do so, we analyzed H3K4me3 distribution in *lsl-1(tm4769)* and *him-17(ok424)* mutant germline. In wildtype germline nuclei, H3K4me3 is broadly distributed on all autosomes (Figure 4 C) (Bean et al., 2004; Li and Kelly, 2011; Xiao et al., 2011). In *lsl-1(tm4769)* and *him-17(ok424)* mutants, the amount of H3K4 trimethylation is globally reduced, particularly in the *him-17(ok424)* mutant germline. This reduced amount of H3K4 trimethylation is associated with an altered distribution, leading to the formation of foci. Furthermore, lack of *lsl-1* activity does not reduce H3K4me3 level uniformly along the gonad. Zone III/IV, corresponding to the transition zone, remains enriched with H3K4 trimethylation. Overall, the X chromosomes remain depleted of H3K4me3 in *lsl-1* and *him-17* mutants (Figure 4 C, white arrows). These results indicate that LSL-1 and HIM-17 are required to maintain normal level of H3K4 trimethylation, which might require the interaction with the COMPASS complex.

### LSL-1 core complex members are required for germ cell fate maintenance

Since LSL-1 is a major activator of the germline genetic program, we wondered if LSL-1 is required for germ cell fate maintenance. To answer this, we investigated the expression of the pan-neuronal marker UNC-119::GFP in *lsl-1(ljm1)* mutant gonads (Ciosk et al., 2006; Maduro and Pilgrim, 1995). *lsl-1(ljm1)* mutant gonads show UNC-119::GFP positive cells in the gonad, while no signal is found in wild-type germlines (Figure 5 A). By taking a closer look, we could distinguish neurite-like structures, supporting the idea that germ cells are reprogrammed into neurons (Figure 5 A, yellow square magnifications and white arrows). Interestingly, we observed two distinct patterns of reprogramming in *lsl-1(ljm1)* mutant gonads. A subset of gonads exhibits GFP positive cells in the proximal part, while others show a more distal or global reprogramming (Figure 5 A). Following this result, we wondered whether LSL-1 interacting partners, members of the core complex, are also involved in germ cell fate maintenance. *xnd-1* was already shown to play a role in germ cell fate maintenance, (Mainpal et al., 2015). Interestingly, *him-17(ok424)* mutant gonads also exhibit UNC-119::GFP positive cells in their gonads, along the entire length, as well as neurite-like structures (Figure 5 A). In *bra-2(ok1171)* mutants, we could detect GFP positive cells in whole worms (Figure 5 A).

**Figure 5.**
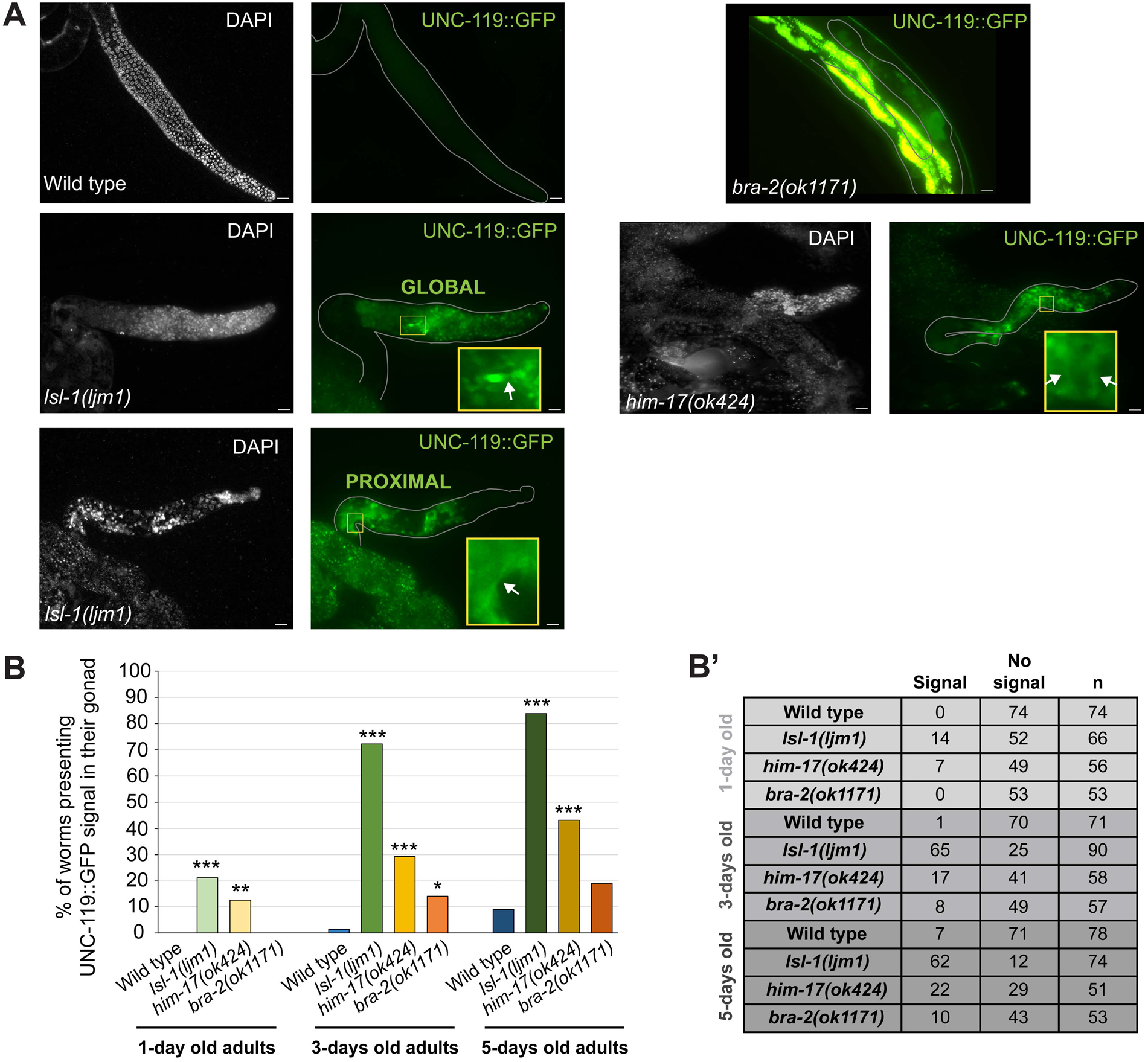
Germ cells lacking LSL-1, HIM-17 or BRA-2 activity are reprogrammed into neuron-like cells. (A) Representative widefield images of 3-days-old adult gonads of wild type, *lsl-1(tm4769)*, *him-17(ok424)* worms expressing the neuronal transgene UNC-119::GFP and stained with DAPI. The *bra-2(ok1171)* mutant expressing UNC-119::GFP is imaged as a whole worm. Yellow squares indicate magnification. Scale bars, 20 µm. (B) Bar plot showing the percentage of gonads presenting UNC-119::GFP positive cells at 1-day-old, 3-days-old and 5-days-old of adults for the indicated genotype. Fisher’s exact test was used to test the significance. One star indicates a p-value < 0.05, two a p-value < 0.01 and three a p-value < 0.001. (B’) Table summarizing the number of gonads with or without UNC-119::GFP signal per strains.

To quantify the extent of germline reprogramming, we monitor the presence of GFP positive cells in mutants at different age of adulthood (Figure 5 B and B’). Mutation in *lsl-1* leads to the highest level of reprogramming, with 72% of the gonads exhibiting reprogramming at 3-days-old adulthood. At the same age, mutants for core members of the LSL-1 complex show 29% for *him-17(ok424)* and 14% for *bra-2(ok1171)*. Altogether, we can conclude that LSL-1 and its core complex partners HIM-17 and BRA-2 play an important role in safeguarding germline fate and preventing reprogramming into neuronal fate.

## Discussion

Our results revealed LSL-1 as a central component of a germline-specific transcriptional protein network, necessary for proper germline development and reproduction, and for safeguarding germline identity. Converging biochemical, genomic, and genetic evidence indicates that LSL-1 interacts with the chromatin factors HIM-17, XND-1, and BRA-2 to form a functional core protein complex that binds to the promoter regions of germline genes and promotes their transcription, likely in collaboration with active chromatin-modifying complexes such as the COMPASS and MOF complexes. We propose that the LSL-1 core complex binds promoter regions of germline-expressed genes within a chromatin environment, where the COMPASS and/or the MOF are present and modulate chromatin organization (Figure 6).

**Figure 6:**
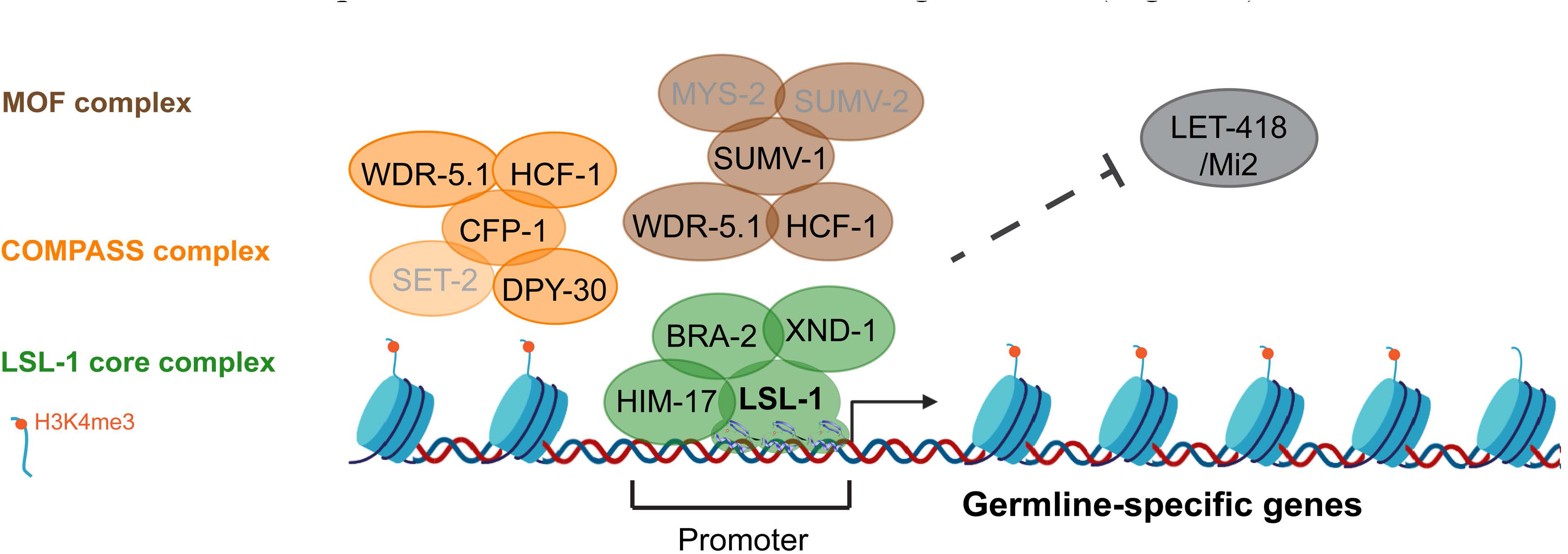
LSL-1 interactors and neighbors activating germline gene transcription. (For details: see text).

CoIP-MS allowed us to identify interactors, that we considered as members of a core complex based on the following observations. The four proteins display overlapping localization patterns on autosomal chromatin in the adult germline, while being excluded from the X chromosomes. ChIP-seq analyses of LSL-1, HIM-17 and XND-1 revealed a substantial overlap in common genomic binding regions, particularly at promoter regions of germline-expressed genes, such as *syp-2* and *nasp-2*. LSL-1 and HIM-17 share a significant overlap of direct target genes, most of which are downregulated. Furthermore, LSL-1 and HIM-17 share common binding motifs that were found as the main binding motifs for LSL-1 and HIM-17 individually (Carelli et al., 2022; Rodriguez-Crespo et al., 2022). In support of our data: XND-1 was also shown to interact *in vitro* with LSL-1 in a Yeast-Two-Hybrid screen and one-to-one interactions between LSL-1, HIM-17, XND-1 and BRA-2 were detected by other labs (Blazickova et al., 2025; Raices et al., 2025; Reece-Hoyes et al., 2013).

The localization analyses further suggest a hierarchical organization within the LSL-1 complex, that might rely on protein domains. Consistent with its main/primary recruiter function, LSL-1 contains three highly conserved C2H2 zinc-finger domains characteristic of the SP/KLF family of transcription factors, and typically enabling DNA binding, as well as two more diverged C2H2 zinc fingers, that could confer other binding specificities, including RNA binding or protein-protein interactions (Bonchuk and Georgiev, 2024; Nabeel-Shah et al., 2024). Similarly, HIM-17 contains six THAP domains, four of them exhibit the canonical residues associated with DNA binding, while two are more divergent and might serve as protein–protein interactions depending on the context (Clouaire et al., 2005; Reddy and Villeneuve, 2004). One possibility is that HIM-17 reinforces chromatin targeting at specific genomic regions already engaged by LSL-1, potentially increasing binding stability or contributing to cooperative DNA recognition. BRA-2 might contribute to the structural integrity or chromatin anchoring of the complex through its MYND-type zinc finger domain, known to mediate protein–protein interactions in chromatin-associated complexes (Spellmon et al., 2015). Lacking known characterized protein domains, XND-1 has been described as a chromatin-associated factor involved in meiotic chromosome regulation and crossover distribution (Mainpal et al., 2015; Wagner et al., 2010). Overall, the biochemical complex composition suggests a role in transcription modulation. However, knowing the role of XND-1, BRA-2 and HIM-17 in crossover regulation and chromosome synapsis, we cannot exclude that the complex is also impacting meiotic functions at the level of chromosome structure (Blazickova et al., 2025; Mainpal et al., 2015; Wagner et al., 2010).

Our proximity labeling data suggest that the LSL-1 core complex further interacts with chromatin modifiers that are associated with transcriptionally active chromatin. The COMPASS is responsible for the methylation of H3K4 (Herbette et al., 2020; Hoe and and Nicholas, 2014). This is consistent with the chromatin landscape observed at LSL-1, HIM-17 and XND-1 binding sites as well as the altered distribution of H3K4me3 in *lsl-1* and *him-17* mutants. The identification of H3K4me3 readers such as SET-9 and SET-26 in the proxisome of LSL-1 further supports a role for of LSL-1 in the balance between deposition, recognition and restriction of H3K4 methylation (Wang et al., 2018). One possibility is that LSL-1 acts by recruiting or stabilizing the COMPASS complex at germline gene promoters, thereby promoting H3K4 methylation and transcriptional activation locally. An alternative possibility is that LSL-1 preferentially binds to promoters already marked by H3K4me3 and contributes to reinforcing this active chromatin state through positive feedback mechanisms. The LSL-1 interaction with the COMPASS is further supported by their involvement in germ cell fate maintenance. Both excessive and insufficient H3K4 methylation have been shown to impact germline identity. Loss of the H3K4 demethylase SPR-5, as well as loss-of-function mutations affecting the COMPASS components SET-2 or WDR-5.1, trigger transdifferentiation of germ cells into somatic cell types, particularly in neurons (Käser-Pébernard et al., 2014; Robert et al., 2015, 2014). These findings indicate that control of H3K4 methylation is essential for maintaining germ cell fate. The partial reduction and redistribution of H3K4me3 observed in *lsl-1* and *him-17* mutants may therefore contribute not only to transcriptional downregulation of germline genes but also to alteration of germ cell fate. We also identified a biochemical and a functional link between LSL-1 and members of the MOF complex. The function of the MOF complex in *C. elegans* is less characterized. However, several studies indicate that a MOF complex plays a role in worm development (reviewed in Hoe and Nicholas, 2014). Interestingly, a crosstalk between the MOF and the COMPASS has been demonstrated biochemically in humans, where hMOF promotes H3K4me2 by the COMPASS complex (Zhao et al., 2013). The interaction of the COMPASS member DPY-30 with the MOF proteins SUMV-1 and SUMV-2 using a yeast-two-hybrid assay is another piece of evidence for a link between the two complexes, in addition to the shared components (Yücel et al., 2014). Altogether, the genetic and the biochemical data, strongly suggest that the LSL-1 core complex functions in association with the COMPASS and the MOF complexes.

Surprisingly, we uncovered no protein of the so-called epigenetic memory of germline gene expression. For the establishment of the proper germline transcriptional program and consequently the survival and development of the PGCs (Primordial Germ Cells), H3K36 methylation and the associated readers and writers, MET-1, MES-4, MRG-1 are key players (Andersen and Horvitz, 2007; Furuhashi et al., 2010; Kelly, 2014; Rechtsteiner et al., 2010). One possible explanation is that these chromatin proteins are acting at the level of the gene body, while LSL-1 is functioning on the promoter (Rechtsteiner et al., 2010).

In conclusion, we propose that an LSL-1 protein complex, in association with chromatin modifiers, is involved in transcription regulation of germline genes, thereby ensuring maintenance of germ cell fate.

## Material and methods

### Worm growth and maintenance

All strains used in this study were kept at 20°C on standard Nematode Growth Media (NGM) agar plates seeded with *Escherichia coli* OP50, according to the standard maintenance conditions (Brenner, 1974) or for the experiments, except when stated otherwise. The wild-type strain used throughout the study was *C. elegans* var. Bristol (N2). Standard genetic crosses were used to generate double mutant strains or strains expressing transgene protein in a mutant background. All the strains used in this study are listed in the Supplementary Methods.

### Molecular cloning and transgenesis

Gibson Assembly was used to generate *pie-1p::egl-13NLS::CeTurboID::mNeongreen::3XFLAG::lsl-1 3’UTR* control strain for the TurboID proximity labeling (Gibson et al., 2009).

The Multisite Gateway Cloning system from Invitrogen (Reece-Hoyes and Walhout, 2018) was used to generate the *bra-2::gfp*, *nasp-2::gfp* and *syp-2::gfp* transgenes, as well as *lsl-1::gfp* transgenes under the control of different 3’UTR.

Integration of the plasmids into the genome was done using the MosSCI insertion (Frøkjær-Jensen et al., 2008) and were inserted on Chromosome II or IV as single copy plasmids. Plasmids and primers used in this study are listed in the Supplementary Methods.

### Protein extraction, complex purification and proximity labeling

The GFP-trapping protocol was adapted from (Pillet et al., 2022) and we used the proximity labeling approach from (Artan et al., 2021). The detailed protocols, as well as mass-spectrometry analysis of the LSL-1 interactome and proxisome, are available in the Supplementary Methods.

### Immunofluorescence and microscopy

Worm gonads were dissected and stained with 4’,6’-diamidino-2-phenylindole hydrochloride (DAPI) or immunostained with H3K4me3 antibody 1:200 (Abcam #ab1012) and subsequent secondary antibody 1:200 (Sigma #T5393) as described by Phillips et al. (2009). At least thirty gonads of each strain from two biological replicates were imaged.

Microscopy pictures were taken through a 100x 1.4 NA objective, at each 0.2 μm thickness intervals, using a pco.edge sCMOS camera connected to a Visitron Visiscope CSU-W1 spinning disk confocal microscope (Nikon Ti/E inverted microscope).

Images were then processed using Fiji ImageJ software as follows: multiple Z-stack projection, ImageJ Stitching plugin (Preibisch et al., 2009), background subtraction, contrast/brightness adjustment, channel merging and scale setting. Final picture arrangement was performed using Adobe Illustrator 2025.

### RNAi treatment

The *E. coli* HT115 bacteria carrying cloned L4440 plasmids expressing double-stranded RNA of the gene of interest are coming from the Ahringer bacterial feeding library (Kamath and Ahringer, 2003). The HT115 bacterial strain carrying the empty L4440 (pPD129.36) cloning vector was used as a control.

P0 L4 larvae were cultured on NGM plates containing 1 mM IPTG and 50 µg/ml ampicillin, seeded with the corresponding HT115 bacteria and grown at 25°C for two or three days to induce RNAi by the feeding technique (Timmons et al., 2001). Their F1 progeny was transferred to a drop of 2% NaN_3_ on a 2% Agar pad and imaged under a fluorescent Zeiss Axioplan 2 microscope.

Images were acquired through 63x or 100x objective with a Zeiss AxioCam color camera driven by AxioVision v4.8.2 software and then processed using Fiji ImageJ software. At least 100 F1 larvae were assessed for each RNAi treatment from at least three different biological replicates. The number of germ cells was determined by counting PGL-1::mCherry positive cells (Figure S4).

Pairwise comparisons of the five-category distributions between RNAi samples were performed using the exact Fisher-Freeman-Halton test. These statistical analyses were conducted on RStudio (version 4.4.1).

### Somatic transdifferentiation

At least 50 worms expressing the neuronal marker *unc-119::gfp* were imaged at one-day-old, three-day-old and five-day-old of adulthood. To quantify the number of worms showing a positive somatic signal, gonads were dissected, fixed according to Phillips et al. (2009) and imaged using a fluorescent Zeiss Axioplan 2 microscope. Images were processed using Fiji ImageJ software.

## Data availablity

The mass spectrometry proteomics data have been deposited to the ProteomeXchange Consortium via the PRIDE (Perez-Riverol et al., 2025) partner repository with the dataset identifier PXD076461.

## Funding

This work was supported by IZCOZ0_198093 (linked to COST Action CA18127 International Nucleome Consortium) and SNSF (Swiss National Science Foundation) Grants 31003A_179395 to CW.

## Acknowledgements

The authors thank L. Schild, C. Déforel and L. Bulliard for excellent technical support; the “model organism Encyclopedia of Regulatory Networks” resource (modERN; http://epic.gs.washington.edu/ modERN/) for ChIP-seq data. Some strains were provided by the Caenorhabditis Genetics Center (CGC; cbs.umn.edu/cgc/home), which is funded by NIH Office of Research Infrastructure Programs (P40 OD010440).

## Notes

### Competing Interest Statement

The authors have declared no competing interest.

