## Supplementary Figures for "The transcription factor LSL-1 interacts with the chromatin factors HIM-17, XND-1 and BRA-2 to promote the germline-specific transcriptional repertoire and to safeguard germ cell fate in *C. elegans*"

Supplementary figures captions

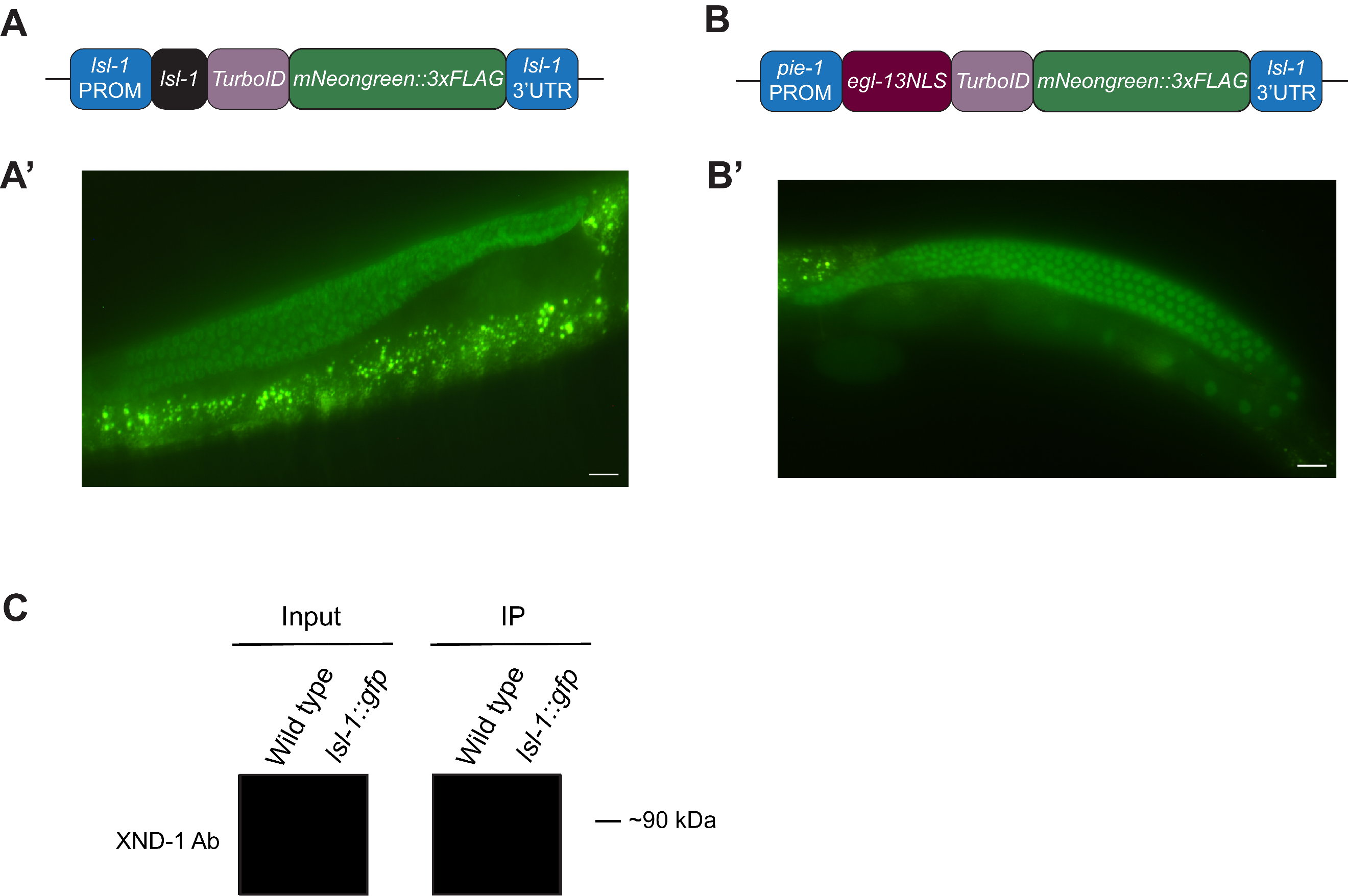

**Figure S1 LSL-1::TurboID and TurboID Ctrl are properly expressed in *C. elegans* germline.** (A and B) Schematic representation of *lsl-1::TurboID* and TurboID control transgenic construction with *pie-1* promoter and *egl-13* nuclear localization signal (NLS). (A’ and B’) Representative widefield pictures of *lsl-1::turboID* CRISPR strain and TurboID control transgenetic strain in one-day-old adult worms. Scale bars, 20 µm. (C) Western blot of wild-type sample and sample of the strain expressing *lsl-1::gfp*, after young adult germline nuclei extraction. Membranes were incubated with anti-XND-1 antibodies for the input (before GFP-trap precipitation) and immunoprecipitated samples. IP, immunoprecipitated.

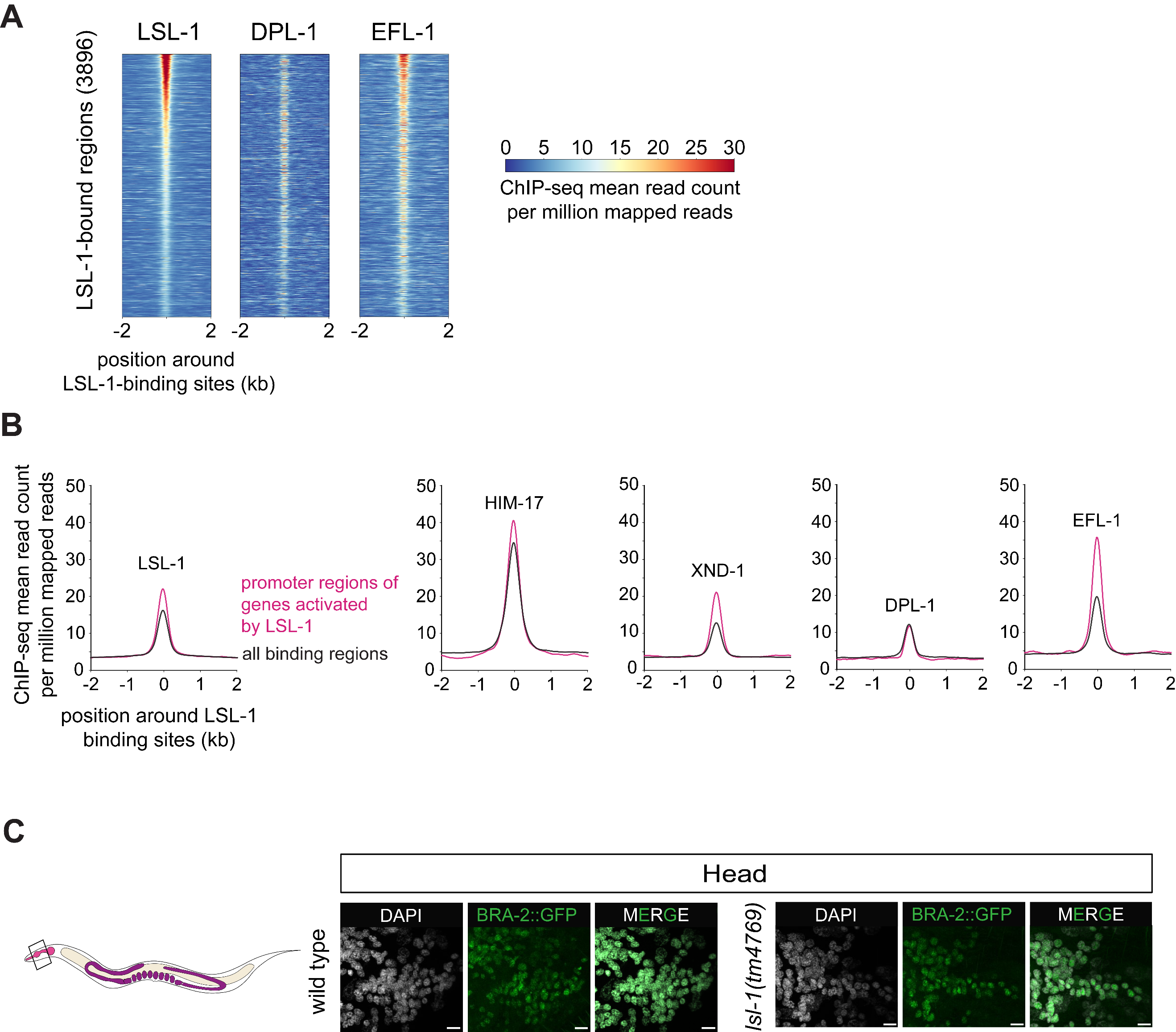

**Figure S2 LSL-1, HIM-17, XND-1 and EFL-1 are enriched at LSL-1 binding sites. (**A) Heatmaps show enrichment of LSL-1, DPL-1 and EFL-1 at the LSL-1-bound regions determined by ChIP-seq peaks (3896). The reads were positioned around the center of the LSL-1 binding sites with 2 kb regions upstream and downstream. (B) Profile plot shows the LSL-1 binding regions enrichment of LSL-1, HIM-17, XND-1, DPL-1 and EFL-1, centered on LSL-1 binding regions (ChIP-seq peaks). The reads were positioned around the center of the LSL-1 binding sites with 2 kb regions upstream and downstream. Black lines show the peak for all LSL-1 binding regions, while pink lines show the LSL-1 binding regions located at the promoter of the genes downregulated in *lsl-1(tm4769)* mutant young adult worms. **(**C) Representative confocal projection images of neuron nuclei of 1-day-old adult stage heads of wild type and *lsl-1(tm4769)* mutant worms, expressing BRA-2::GFP. Scale bars, 5 µm.

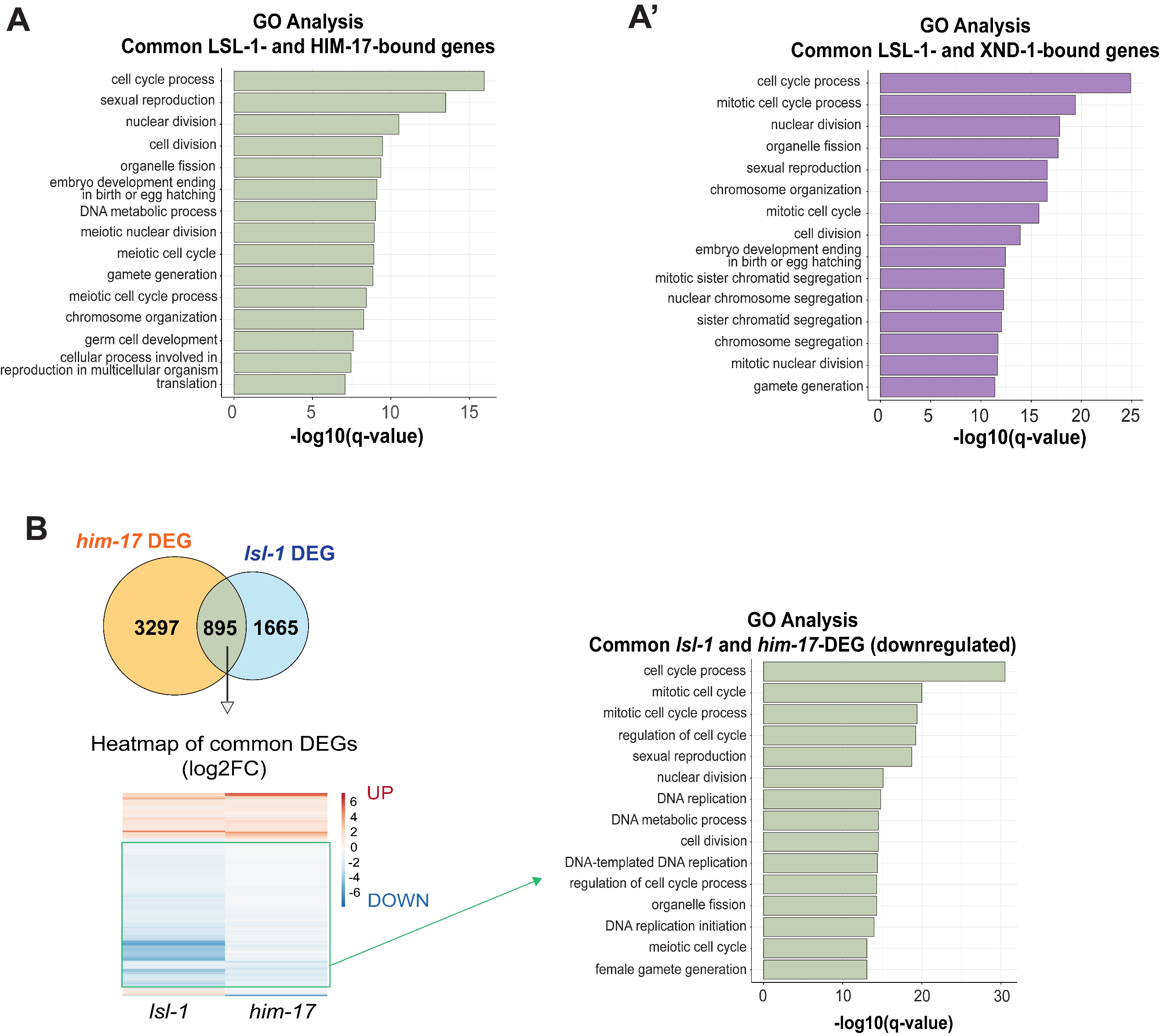

**Figure S3 LSL-1, HIM-17 and XND-1 common bound genes are involved in germline processes. LSL-1 and HIM-17 promote the expression of genes involved in germline processes.** (A and A’) Bar plots representing the Gene Ontology enrichment analysis of the common gene lists of LSL-1 vs HIM-17, or XND-1, ChIP-seq datasets. GO terms with a q value < 0.05 were considered significantly enriched. (B) Overlap between HIM-17 (orange) and LSL-1 (blue) DEGs from published transcriptomics data. Heatmap illustrates the log2Foldchange of DEGs in both *lsl-1* and *him-17* mutants, where red indicates upregulated genes, and blue downregulated ones. Bar plots represent the Gene Ontology enrichment analysis of the common gene lists. GO terms with a q value < 0.05 were considered significantly enriched.

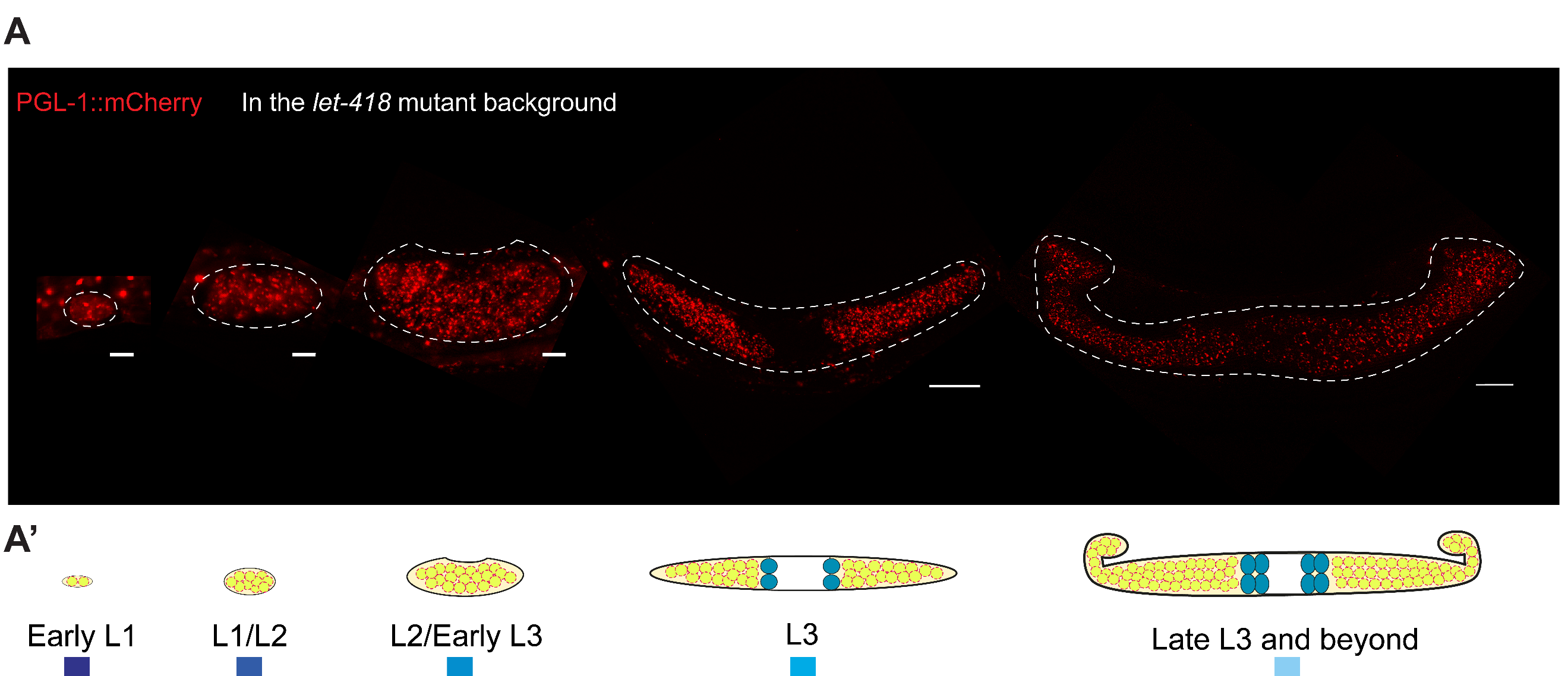

**Figure S4 PGL-1::mCherry is distributed around germline nuclei during germ cell development.** (A) Representative widefield images of germ cells of 3-day-old *let-418(n3536)* mutant progeny expressing PGL-1::mCherry and treated with RNAi, at each developmental stage category according to the suppression of *let-418* mutant phenotype. Scale bars, 5 µm for Early L1, L1/L2 and L2/Early L3 pictures and 20 µm for L3 and Late L3 and beyond pictures. (A’) Drawing of the corresponding developmental stage category.
