## Supplementary Methods for "The transcription factor LSL-1 interacts with the chromatin factors HIM-17, XND-1 and BRA-2 to promote the germline-specific transcriptional repertoire and to safeguard germ cell fate in *C. elegans*"

### Supplementary materials and methods

##### List of strains used and generated in this study

N2 (Bristol)	Wildtype

AV280	*unc-119(e2498) III; him-17(ok424) V; meIs5 [him-17::GFP + unc-119(+)]*

CER414	*pgl-1(cer70[pgl-1::mcherry]) IV*

CWC19	*lsl-1(ljm1) I/tmC18[dpy-5(tmls1200)] I*

CWC22	*lsl-1(tm4769) I/tmC18[dpy-5(tmls1200)] I*

CWC37	*lsl-1(ljm1) I/tmC18[dpy-5(tmls1200)] I; edls6 [unc-119::gfp + rol-6(su1006)] IV*

CWC52	*pgl-1(cer70[pgl-1::mcherry]) IV; let-418(n3536) V*

CWC61	*lsl-1(syb3772[lsl-1::GFP]) I*

CWC79	*lsl-1(syb4742[lsl-1::CeTurboID::mNeongreen::3XFLAG]) I* (backcrossed 5x)

CWC91	*lsl-1(ljm1) I/tmC18[dpy-5(tmls1200)] I; mgIs25[unc-97::gfp]*

CWC92	*ljmSi16[pie-1p::egl-13NLS::TurboIDmNeonGreen::lsl-1 3’UTR] II*

CWC99	*ljmSi18[bra-2p::bra-2::gfp::bra-2 3’UTR] IV*

CWC104	*ljmSi19[lsl-1p::lsl-1::gfp::lsl-1 3’UTR] II*

CWC105	*ljmSi20[lsl-1p::lsl-1::gfp::tbb-2 3’UTR] II*

CWC106	*lsl-1(tm4769) I/tmC18[dpy-5(tmls1200)] I; meIs5 [him-17::GFP + unc-119(+)]*

CWC120	*lsl-1(tm4769) I/tmC18[dpy-5(tmls1200)]I; pgl-1(cer70[pgl-1::mcherry]) IV; let-418(n3536) V*

CWC123	*lsl-1(tm4769) I/tmC18[dpy-5(tmls1200)]I; edls6 [unc-119::gfp + rol-6(su1006)] IV*

CWC126	*bra-2(ok1171) III; ljmSi18[bra-2p::bra-2::gfp::bra-2 3’UTR] IV*

CWC127	*bra-2(ok1171) III/sC1(s2023) [dpy-1(s2170) umnIs41] III*

CWC136	*lsl-1(syb3772[lsl-1::GFP]) I; bra-2(ok1171) III/sC1(s2023) [dpy-1(s2170) umnIs41] III*

CWC137	*lsl-1(tm4769) I/tmC18[dpy-5(tmls1200)] I; ljmSi18[bra-2p::bra-2::gfp::bra-2 3UTR] IV*

CWC139	*lsl-1(tm4769) I/tmC18[dpy-5(tmls1200)] I; eaIs6 [3xFLAG::GFP::SBP::xnd-1+unc-119(+) transgene]*

CWC147	*lsl-1(syb3772[lsl-1::GFP]) I; xnd-1(ok709) III*

CWC165	*lsl-1(tm4769) I; ljmSi20[lsl-1p::lsl-1::gfp::tbb-2 3’UTR] II*

CWC174	*him-17(ok424) V/nT1[q1Iss1] V*

CWC177	*lsl-1(syb3772[lsl-1::GFP]) I; him-17(ok424) V/nT1[q1Iss1] V*

CWC179	*edls6 [unc-119::gfp + rol-6(su1006)] IV ; him-17(ok424) V/nT1[q1Iss1] V*

CWC188	*bra-2(ok1171) III/sC1(s2023) [dpy-1(s2170) umnIs41] III; edls6 [unc-119::gfp + rol-6(su1006)] IV*

CWC189	*ljmSi30[syp-2::syp-2::gfp::syp-2 3’UTR] IV*

CWC190	*ljmSi31[nasp-2::nasp-2::gfp::nasp-2 3’UTR] IV*

CWC191	*ljmSi30[syp-2::syp-2::gfp::syp-2 3’UTR] IV ; him-17(ok424) V/nT1[q1Iss1] V*

CWC192	*ljmSi31[nasp-2::nasp-2::gfp::nasp-2 3’UTR] IV ; him-17(ok424) V/nT1[q1Iss1] V*

CWC193	*lsl-1(tm4769) I/tmC18[dpy-5(tmls1200)] I; ljmSi30[syp-2::syp-2::gfp::syp-2 3’UTR] IV*

CWC194	*lsl-1(tm4769) I/tmC18[dpy-5(tmls1200)] I; ljmSi31[nasp-2::nasp-2::gfp::nasp-2 3’UTR] IV*

DP132	*edls6 [unc-119::gfp + rol-6(su1006)] IV*

EG8885	*unc-119(ed3) III; oxTi933 [eft-3p::GFP::2xNLS::tbb-2 3'UTR + Cbr-unc-119(+)] V*

OH122	*mgIs25[unc-97::gfp]*

PH4742	*lsl-1(syb4742[lsl-1::CeTurboID::mNeongreen::3XFLAG]) I*

QP922	*unc-119 xnd-1(ok709) III; eaIs6 [3xFLAG::GFP::SBP::xnd-1+unc-119(+) transgene]*

RB869	*xnd-1(ok709) III*

The CWC22: *lsl-1(tm4769) I/tmC18[dpy-5(tmls1200)] I* was generated from FX04769: *lsl-1(tm4769) I/(+) I*, a strain created by the National Bioresource Project, *C. elegans* Gene Knockout Consortium; Tokyo, Japan (The C. elegans Deletion Mutant Consortium, 2012).

The QP922: *unc-119 xnd-1(ok709) III; eaIs6 [3xFLAG::GFP::SBP::XND-1+unc-119(+) transgene]* strain was kindly provided by Judith L. Yanowitz (Magee-Womens Research Institute of the University of Pittsburgh, Pittsburgh PA, USA), and is a strain generated by Rana Mainpal (Mainpal et al., 2015; Mainpal and Yanowitz, 2016).

The CER414: *pgl-1 (cer70[pgl-1::mcherry]) IV* strain was kindly provided by Julián Cerón (*C. elegans* Core Facility-IDIBELL; Barcelona, Spain).

CWC61: *lsl-1(syb3772[lsl-1::GFP]) I* was generated by backcrossing PHX3772: *lsl-1(syb3772)* to N2 wild-type background four times. SunyBiotech Company (Fuzhou, China) generated this strain for us by knocking-in GFP sequence at the C-terminal of the endogenous *lsl-1* gene sequence, thanks to CRISPR-Cas9 engineering tools.

The same company generated for us PH4742: *lsl-1(syb4742[lsl-1::CeTurboID::mNeongreen::3XFLAG]) I* strain. The *CeTurboID::mNeongreen::3XFLAG* sequence was designed by Murat Artan from Mario de Bono lab (Artan et al., 2021) and was knocked in at the C-terminal of the endogenous *lsl-1* gene sequence thanks to CRISPR-Cas9 engineering tools. The resulting strain was then backcrossed five times, and its construct expression was verified by fluorescent widefield microscopy (Figure S1).

The remaining strains used in this study were either obtained from the *Caenorhabditis* Genetics Center (CGC, University of Minnesota; Minneapolis MN, USA), funded by the National Institutes of Health (NIH; Bethesda MD, USA), or generated by crosses.

The *CeTurboID::mNeongreen::3XFLAG* sequence used to generate the PH4742 strain is provided here:

GlyLinker-TurboID-mNeongreen-3xFLAG

GGAGGTGGTGGATCAGGCTCGGGAGGTCGAGGCTCAGGATCCGGTTCCGGCTCCGGCTCTGGTTCCGGTTCGGGTTCCGGTTCTGGAAAGGATAACACCGTTCCACTTAAGCTTATCGCCCTTCTTGCCAACGGAGAATTCCACTCTGGAGAGCAACTTGGAGAGACTCTTGGAATGTCCCGTGCTGCCATCAACAAGCATATCCAAACCCTTCGTGATTGGGGAGTTGATGTTTTCACTGTTCCAGGAAAGgtaagtttaaacatatatatactaactaaccctgattatttaaattttcagGGATACTCCCTTCCAGAGCCAATCCCACTTCTTAACGCCAAGCAAATCCTTGGACAACTTGATGGAGGATCCGTCGCTGTCCTTCCAGTTGTTGATTCCACCAACCAATACCTTCTTGACCGTATCGGAGAGCTTAAGTCTGGAGACGCCTGCATCGCTGAGTACCAACAAGCTGGACGCGGATCTCGCGGACGCAAGTGGTTCTCCCCATTCGGAGCCAACCTTTACCTTTCTATGTTCTGGCGTCTTAAGCGTGGACCAGCTGCTATCGGACTTGGACCAGTTATCGGAATCGTTATGGCTGAGGCCCTTCGTAAGCTTGGAGCTGATAAGgtaagtttaaacagttcggtactaactaaccatacatatttaaattttcagGTTCGTGTTAAGTGGCCAAACGATCTTTACCTTCAAGACCGTAAGCTTGCTGGAATCCTTGTCGAGCTTGCTGGAATCACCGGAGACGCCGCTCAAATCGTTATCGGAGCTGGAATCAACGTTGCCATGCGTCGTGTTGAGGAGTCTGTTGTTAACCAAGGATGGATCACTCTTCAAGAGGCTGGAATCAACCTTGATCGTAACACCCTTGCTGCCACCCTTATCCGTGAGCTTCGTGCTGCCCTTGAGCTTTTCGAGCAAGAGGGACTTGCCCCATACCTTCCACGCTGGGAGAAGCTTGACAACTTCATCAACCGCCCAGTTAAGCTTATCATCGGAGATAAGGAAATCTTCGGAATCTCTCGCGGAATCGACAAGCAAGGAGCTCTTCTTCTTGAGCAAGATGGAGTCATTAAGCCATGGATGGGAGGAGAGATTTCCCTTCGTTCCGCTGAGAAGGCCGGAGGAGAACAGAAGCTTATAAGTGAGGAGGACCTGGGATCCGCTGGATCCGCTGCTGGATCCGGTGAGTTCATGGTGTCGAAGGGAGAAGAGGATAACATGGCTTCACTCCCAGCTACACACGAACTCCACATCTTCGGATCGATCAACGGAGTGGATTTCGATATGGTCGGACAAGgtaagtttaaacatatatatactaactaaccctgattatttaaattttcagGAACTGGAAACCCAAACGATGGATACGAGGAACTCAACCTCAAGTCGACAAAGGGAGATCTGCAATTCTCGCCATGGATTCTCGTGCCACACATCGGATACGGATTCCACCAATACCTCCCATACCCAGgtaagtttaaactgagttctactaactaacgagtaatatttaaattttcagATGGAATGTCACCATTCCAAGCTGCCATGGTGGATGGATCGGGATACCAAGTTCACCGAACAATGCAATTCGAGGATGGAGCCTCGCTCACAGTGAACTACCGATACACATACGAGGGATCGCACATCAAGgtaagtttaaacagttcggtactaactaaccatacatatttaaattttcagGGAGAGGCTCAAGTTAAGGGAACAGGATTCCCAGCTGATGGACCAGTGATGACAAACTCACTCACAGCTGCTGATTGGTGCCGATCGAAAAAGACATACCCAAATGATAAGACAATCATCTCGACATTCAAGTGGTCGTACACTACTGGAAACGGAAAGCGATACCGATCGACAGCCCGAACAACATACACATTCGCTAAGCCAATGGCCGCCAACTACCTCAAGgtaagtttaaacatgattttactaactaactaatctgatttaaattttcagAATCAACCAATGTACGTGTTCCGAAAGACAGAACTCAAGCACTCAAAGACAGAGCTGAACTTCAAAGAGTGGCAAAAGGCCTTCACAGATGTGATGGGAATGGATGAACTCTACAAGGACTACAAAGACCATGACGGTGATTATAAAGATCATGACATCGATTACAAGGATGACGATGACAAG

##### Plasmids used in this study

The new plasmids generated for this study are listed in the following table with the information to generate them:

| **Desired sequence** | **Primer containing att recombination sites** | **Entry clone** | **Plasmid name** |
| --- | --- | --- | --- |
| *bra-2* promoter+ORF | pCW 259: GGGG ACA ACT TTG TAT AGA AAA GTT G ATCAGCGACAGGATCAAGCA  pCW260: GGGG AC TGC TTT TTT GTA CAA ACT TGT TTGTTGTTGTGGCGGCATTG | pDONRP4-P1R | pCW116 |
| *bra-2* 3’UTR | pCW 262: GGGG ACA GCT TTC TTG TAC AAA GTG GGA TAATTTTCTTTCGTTTTAACTCTCTAATCATCA  pCW263: GGGG AC AAC TTT GTA TAA TAA AGT TG TTTTCCGCAGATGAAGACGC | pDONRP2R-P3 | pCW117 |
| *nasp-2* promoter | PrCW318: GGGG ACA ACT TTG TAT AGA AAA GTT G ATATTTCCGCAAGCAGGGTG  PrCW313: GGGG AC TGC TTT TTT GTA CAA ACT TGT CAT TATGAAGTTAGTAGTCTGAAAATAATAG | pDONRP4-P1R | pCW106 |
| *nasp-2* ORF | prCW314: GGGG ACA AGT TTG TAC AAA AAA GCA GGC TCAAAAATGAGCGCTTTTGCTGATTCTAC  prCW315: GGGG AC CAC TTT GTA CAA GAA AGC TGG GT TCA ACTAGTGCG GCCGCCTTCGTTGATTGGTGTTGGCTC | pDONR201 | pCW109 |
| *nasp-2* 3’UTR | prCW316: GGGG ACA GCT TTC TTG TAC AAA GTG GGA TAA TTCGTCGTCTTTCTCTCCAATTTC  prCW317: GGGG AC AAC TTT GTA TAA TAA AGT TG ATTTAATAAAATCACAAGTCGCCTTC | pDONRP2R-P3 | pCW110 |
| *syp-2* promoter | prCW319: GGGG ACA ACT TTG TAT AGA AAA GTT G GGAACGTTCAACAATCCAAGC prCW311: GGGG AC TGC TTT TTT GTA CAA ACT TGT CATTCAAAAATTCTCAGAAGCGAAGTTT | pDONRP4-P1R | pCW115 |
| *syp-2* ORF | prCW307: GGGG ACA AGT TTG TAC AAA AAA GCA GGC TCA AAAATGAATTCTGCCCATCGATCTTCG  prCW308: GGGG AC CAC TTT GTA CAA GAA AGC TGG GT TCA ACTAGTGCGGCCGCCTAACTTGTCAGCCCACGGCTCG | pDONR201 | pCW107 |
| *syp-2* 3’UTR | prCW309: GGGG ACA GCT TTC TTG TAC AAA GTG GGA TAA ATCATCTGTTGTATTCAATTTCCTTG  prCW310: GGGG AC AAC TTT GTA TAA TAA AGT TG GTTTCATTTTGTCCTGTGTCCTA | pDONRP2R-P3 | pCW108 |
| *nasp-2* ORF+gfp | – | pDONR201 | pCW120 |
| *syp-2* ORF+gfp | – | pDONR201 | pCW102 |

LR reactions were performed to fuse a combination of the different entry clones into the pCFJ150 backbone using LR clonase II (Invitrogen; Carlsbad CA, USA) to generate the following constructions:

pCW99 : *bra-2p::bra-2::gfp::bra-2 3’UTR* from pCW116, pCM1.53, pCW117 and pCFJ150.

pCW111 : *nasp-2p::nasp-2::gfp::nasp-2 3’UTR* from pCW106, pCW120, pCW110 and pCFJ150.

pCW105: *syp-2p::syp-2::gfp::syp-2 3’UTR* from pCW115, pCW102, pCW108 and pCFJ150.

pCW118: *lsl-1p::lsl-1::gfp::lsl-1 3’UTR* from pCW43, pCW54, pCW45 and pCFJ150.

pCW119: *lsl-1p::lsl-1::gfp::tbb-2 3’UTR* from pCW43, pCW54, pCM1.36 and pCFJ150.

##### ChIP-seq data and RNA-seq data *in silico* analysis

We used ChIP sequencing data available from the <https://www.encodeproject.org/> website, containing a large amount of data collected by the model organism Encyclopedia of Regulatory Networks (modERN) consortium (Kudron et al., 2018; The ENCODE Project Consortium et al., 2012). In addition, other ChIP sequencing data used were found in publications or from the model organism encyclopedia of DNA Elements (modENCODE) project. They are all listed in the following Table X according to their reference:

| **Target protein or histone mark** | **GEO accession number** | **Reference** |
| --- | --- | --- |
| LSL-1::TY1::EGFP::3xFLAG | GSE258740 | (Kudron et al., 2018)  (The ENCODE Project Consortium, 2012) |
| XND-1::EGFP::3xFLAG | GSE257106 | (Kudron et al., 2018)  (The ENCODE Project Consortium, 2012) |
| HIM-17 | GSE192540 | (Carelli et al., 2022) |
| EFL-1::EGFP::3xFLAG | GSE258267 | (Kudron et al., 2018)  (The ENCODE Project Consortium, 2012) |
| DPL-1::EGFP::3xFLAG | GSE258502 | (Kudron et al., 2018)  (The ENCODE Project Consortium, 2012) |
| H3K4me3 | GSE114440 | (Jänes et al., 2018) |
| H3K36me3 | GSE147401 | (McManus et al., 2021) |
| H3K79me2 | GSE49742 | modEncode_5169 project of Jason Lieb |
| H3K9me2 | GSE87522 | (McMurchy et al., 2017) |
| H3K9me3 | GSE87522 | (McMurchy et al., 2017) |
| H3K27me3 | GSE174652 | (Zaghet et al., 2021) |

All bioinformatic analyses were performed on the Galaxy web platform using the public server at https://usegalaxy.org (Afgan et al., 2016) and RStudio (version 4.4.1).

These ChIP-seq experiments were made in young adult wildtype worms (for HIM-17 ChIP and all histone marks) or young adult worms expressing either LSL-1::TY1::EGFP::3xFLAG (wgls720), XND-1::EGFP::3xFLAG, DPL-1::EGFP::3xFLAG and EFL-1::EGFP::3xFLAG transgenes. The proteins of interest were immunoprecipitated with anti-HIM-17 antibody or anti-GFP antibody, respectively, for the proteins, and antibodies against the corresponding histone marks for them. Resulting sequencing data were aligned to the C. elegans reference genome WS245 using the Burrows-Wheeler aligner (BWA) (Li and Durbin, 2009). Next, peak calling was performed using the ChIP-seq pipeline SPP (Kharchenko et al., 2008) or the MACS2 tool on the Galaxy web platform using the public server at https://usegalaxy.org (Afgan et al., 2016) on the regions significantly enriched in aligned reads. The final peaks were filtered by those with an IDR above 0.1% (Landt et al., 2012). The .bed file available on the modern website was annotated through Galaxy using the *ChIPseeker R/bioconductor package* (version 1.18.0) (Yu et al., 2015).

ChIP-seq data of the histone marks were obtained from young adult worms, except H3K36me3 from dissected young adult gonads and H3K79me2 from early embryos. To analyze the chromatin environment around LSL-1, HIM-17 and XND-1 binding sites, ChIP-seq data from active and repressive chromatin marks – H3K4me3, H3K36me3, H3K79me2, H3K9me2, H3K9me3 and H3K27me3 – were used. Their fastaq files were filtered by quality, also mapped to the *C. elegans* reference genome W245 with BWA and sorted (Samtool) (Li and Durbin, 2009). After removing duplicates, reads were (RPKM)-normalized using the *bamCoverage* tool and plotted using *computeMatrix* and *plotProfile* tools from the deepTools2 function (deeptools v3.3.2, samtools v1.9) (Ramírez et al., 2016).

The resulting lists of genes were LSL-1, HIM-17 and XND-1 bind were updated for their name and WormbaseID using the SimpleMine and Gene Name Sanitizer tools available on https://wormbase.org/tools. They were then compared together using RStudio (version 4.4.1), plotted into Venn diagrams, and common genes were analyzed further for Gene Ontology (GO) enrichment to identify overrepresented biological processes among them. GO enrichment analysis was conducted using the *clusterProfiler* R package (version 4.14.6). WormBase gene identifiers (WormbaseID) were used as input, and enrichment was performed against the *Caenorhabditis elegans* genome using the *org.Ce.eg.db* annotation package. Only Biological Process (BP) ontology was considered in this study. Enrichment significance was assessed using a hypergeometric test, and resulting *p*-values were corrected for multiple testing using the Benjamini–Hochberg false discovery rate (FDR) method. GO terms with an adjusted *p*-value (q-value) < 0.05 were considered significantly enriched.

Enrichment results were visualized using bar plots generated with *ggplot2*. Bar plots represent the statistical significance expressed as −log10(q-value) associated with each GO term. For visualization purposes, the top 15 enriched GO terms were selected based on adjusted *p*-value.

Similar *in silico* analyses with RStudio (version 4.4.1) were performed on the gene lists obtained for differentially-expressed genes (DEG) in *lsl-1* and *him-17* mutant worms through available transcriptomics data (Carelli et al., 2022; Rodriguez-Crespo et al., 2022). For these analyses, we chose to keep all the DEGs with log2 Fold Change above or below 0.

Thanks to the gene lists of ChIP-seq and transcriptomics data for LSL-1 and HIM-17, a list of direct target genes (bound at the promoter and DEG) and a list of indirect target genes (not bound but DEG) were determined, compared and analyzed for Gene Ontology (GO) enrichment. From the final common list of direct target genes between LSL-1 and HIM-17, a 2-kb length genomic region that is upstream of the annotated coding gene start site was extracted for each common genes using the coding_gene_flank option of the *biomaRt* package on RStudio, which query the Ensembl BioMart database for *Caenorhabditis elegans* (dataset: *celegans_gene_ensembl*), as promoter sequences. They were then converted into a *DNAStringSet* object using the *Biostrings* package, and named according to their corresponding WormBase gene ID. Finally, all promoter sequences were exported in FASTA format to generate a single file suitable for downstream computational analyses, such as motif discovery.

These promoter sequences in FASTA format were analyzed for *de novo* motif enrichment using MEME (version 5.5.9) web-based tool (<https://meme-suite.org/meme/>) (Bailey et al., 2015; Bailey and Elkan, 1994). MEME was run in DNA classic mode using the Zero or One Occurrence Per Sequence (ZOOPS) model. The maximal number of motifs to identify was set to 10; motif widths were allowed to vary between 5 and 12 nucleotides. Motifs were identified and ranked according to their E-values.

##### Protein extraction and complex purification using the GFP-trapping system

Wild-type worms, worms expressing a nuclear GFP (*eft-3p::GFP::2xNLS::tbb-2 3'UTR*) and worms expressing a CRISPR-generated *lsl-1::gfp* knock-in at the native locus were amplified and synchronized by standard bleaching, and grown at 20°C on 15cm NGM plates seeded with OP50 *E. Coli* until one-day-old adult stage. They were collected, washed with M9 and a last wash with MilliQ water, pelleted into a packed pellet of 400 µl and flash-frozen in liquid nitrogen.

The remaining steps of the procedure were performed on ice or at 4°C. Worms were broken into the same volume of the pellet of modified TNN-150 lysis buffer using a micropestle (weak homogenizer), and sonicated 2x 10 seconds at 70% amplitude with 30 seconds cool down in between with the Q125 Sonicator® (Qsonica). Cell lysates were clarified by spinning at 5’000 rpm for 10 minutes at 4°C, and their supernatant was collected into new 1.5 ml tubes. Their protein concentration was measured using the BCA^TM^ Protein Assay Kit – Reducing Agent Compatible (Thermo Scientific).

The GFP-trapping protocol was adapted from (Pillet et al., 2022). GFP-Trap® Magnetic Agarose beads (ChromoTek) were washed with cold modified TNN-150 lysis buffer without NP-40, DTT, PMSF and protease inhibitor cocktail, and blocked at 4°C during 1-2 hours on a rotating wheel with 3% BSA in modified TNN-150 without NP-40, DTT, PMSF, and protease inhibitor cocktail. 20 µl of blocked GFP-Trap bead slurry were incubated with 5-10 mg of protein from the cell lysate for 2 hours at 4°C on a rotating wheel. Beads were then washed 3x with modified TNN-150 buffer without NP-40, DTT, PMSF and protease inhibitor cocktail. Then, beads were either boiled for 5 minutes at 95°C in 50 µl of 3x SDS sample buffer to elute the proteins for Western blotting or kept at -80°C for on-beads digestion for mass spectrometry analysis.

##### Proximity-labeling, extraction and precipitation of biotinylated proteins - TurboID

The protocol was adapted from (Artan et al., 2021).

Wild-type worms, worms expressing *pie-1p::egl-13NLS::TurboIDmNeonGreen::lsl-1 3’UTR* and worms expressing a CRISPR-generated *lsl-1::TurboIDmNeonGreen* knock-in at the native locus were amplified and synchronized by standard bleaching and grown at 20°C on 15cm NGM plates seeded with biotin auxotroph MG1655 *E. Coli* until the young adult stage. They were collected and washed with M9 until the supernatant was cleared of bacteria. They were then incubated in M9 supplemented with 1 mM biotin and MG1655 bacteria for 2 hours at 22°C on a rotator at 11-12 rpm, before being washed again with M9 to get rid of the bacteria. The collected worms were finally washed with MilliQ water once, pelleted into a packed pellet of 400 µl, and flash frozen in liquid nitrogen before being kept at -70°C.

Thawed worm pellets were added with one volume of RIPA buffer (50 mM Tris-HCl pH 7.5, 150 mM NaCl, 1% NP-40, 0.5% sodium deoxycholate, 0.1% SDS, 1 mM EDTA, 1 mM PMSF, one tablet of cOmplete^™^, EDTA-free Protease Inhibitor Cocktail (Roche) for 50 ml buffer) and broken with a micropestle (weak homogenizer). Broken worms were incubated at 90°C for 5 minutes and then sonicated twice for 30 seconds at 70% amplitude with 60 seconds cool down in between using the Q125 Sonicator® (Qsonica). Samples were cooled down to room temperature for the addition of urea to a final concentration of 2 M. Cell lysates were clarified by spinning at max speed for 45 minutes at 22°C, and their supernatant was collected into a 1.5 ml tube. Lysates were desalted using Zeba spin desalting columns (7k MWCO) to get rid of free biotin and their protein concentration was measured using the BCA^TM^ Protein Assay Kit – Reducing Agent Compatible (Thermo Scientific). Dynabeads MyOne Streptavidin C1 (Invitrogen) were washed with RIPA lysis buffer without NP-40, sodium deoxycholate, SDS, PMSF and protease inhibitor cocktail. Equilibrated bead slurry was incubated with 5-10 mg protein from the cell lysate overnight under rotation at room temperature. Beads were washed 3x with 2% SDS buffer (150 mM NaCl, 1 mM EDTA, 2% SDS, 50 mM Tris-HCl pH 7.5), 3x with KCl wash buffer (1 M KCl, 1 mM EDTA, 50 mM Tris-HCl pH 7.5) and 2x with TBS buffer (150 mM NaCl, 50 mM Tris-HCl). Then, beads were either boiled twice for 5 minutes at 95°C in 40 µl of 4x NuPAGE LDS sample buffer with 10x NuPAGE sample reducing agent mix saturated with free biotin to elute the proteins for Western blotting or kept at -80°C for on-beads digestion for mass spectrometry analysis.

##### Immunoblotting – Western blotting

Western blots were performed according to standard protocol. Membranes were blocked with 5% Blotting Grade Blocker Non-Fat Dry Milk (Bio-Rad) in 1x PBST, and antibodies were diluted in the same buffer. SuperSignal® West Pico Chemiluminescent Substrate (Thermo Scientific) and the LI-COR Odyssey® FC machine were used to detect Horseradish Peroxidase (HRP) used as detectable signal in antibodies for Western blots, on the membrane.

GFP was detected with anti-GFP from mouse IgG_1_κ (monoclonal clones 7.1 and 13.1) (Roche #11814460001) primary antibody at a dilution of 1:1’000 and a Peroxidase AffiniPure® Goat Anti-Mouse IgG (H+L) (Jackson Immuno Research #115-035-003) secondary antibody at a dilution of 1:5’000. XND-1 was detected with guinea pig anti-XND-1 primary antibody (gift from Judith Yanowitz lab) at a dilution of 1:1’000 and an anti-Guinea Pig IgG (whole molecule)−Peroxidase antibody produced in goat secondary antibody (Sigma Aldrich® #A7289) at a dilution of 1:2’000. The rabbit polyclonal Anti-Histone H3 primary antibody (Abcam #ab1791) at a dilution of 1:1’000 was used as a standard loading control (for nuclear proteins) and an anti-Rabbit IgG (whole molecule)–Peroxidase antibody produced in goat secondary antibody (Sigma Aldrich® #A6154) at a dilution of 1:5’000. Primary antibodies were incubated overnight at 4°C under shaking, while secondary antibodies were incubated for 1 hour at room temperature under shaking.

Biotinylated proteins were detected with Streptavidin-HRP (Cell Signaling® #3999) at a dilution of 1:2’000. It was incubated for 2 hours at room temperature under shaking.

##### Mass spectrometry – sample preparation

Protein samples for all experiments were processed for mass spectrometry analysis using the on-beads digestion protocol provided by the Proteomics Platform (MAPP) of the University of Fribourg.

Beads carrying protein samples were reduced with 1 mM DTT and subsequently alkylated with 5.5 mM IAA, before being transferred on a 10 kD MWCO HY filtrate tube (Vivacon^®^ 500 #VN01H02, Sartorius) for washes with 8 M urea and ABC buffer (100 mM Ammonium bicarbonate, pH 7.5). Washed proteins were digested overnight at 37°C under agitation with 0.2 µg/µl Sequencing Grade Modified Trypsin (Promega). Digested peptides were eluted the next day and washed with ABC buffer. 50% TFA was added to get the pH < 2 to inactivate the trypsin. Eluted peptides were then washed and concentrated using a StageTip procedure as follows (to desalting, purifying and conditioning of peptides). Homemade C18 StageTip (with x20 membrane empore C18 47 mm, Supelco) was equilibrated first with buffer B (80% acetonitrile, 0.5% acetic acid, 20% H_2_O), and then twice with buffer A (0.5% acetic acid in H_2_O). Peptide samples were loaded on them, washed with buffer A and eluted with buffer B. Organic solvents were evaporated using speed-vacuum to decrease the volume to 5 µl. Samples were top up until 20 µl with buffer A*/buffer A (30% A* – 3% acetonitrile, 0.3% TFA –, 70% A, < 1% acetonitrile) and stored at -80°C until their loading on the mass spectrometer.

##### Mass spectrometry analysis

LC-MS/MS measurements were performed on a QExactive HFX mass spectrometer coupled to a Vanquish NEO HPLC (all Thermo Scientific). Peptides were separated on a fused silica HPLC-column tip (I.D. 75 μm, New Objective, self-packed with ReproSil-Pur 120 C18-AQ, 1.9 μm (Dr. Maisch) to a length of 20 cm) using a gradient of A (0.1% formic acid in water) and B (0.1% formic acid in 80% acetonitrile in water). Mass spectrometer was operated in the data-dependent mode; after each MS scan (mass range m/z = 370 – 1750; resolution: 120’000) a maximum of twelve MS/MS scans were performed using an isolation window of 1.6, a normalized collision energy of 28%, a target AGC of 4500 and a resolution of 30’000. Spray voltage was set to 2.3 kV and the ion-transfer tube temperature to 250°C; no sheath and auxiliary gas were used.

MS raw files were analyzed using MaxQuant (version 2.0.1.0) (Cox and Mann, 2008) using a UniProt full-length *C. elegans* database and common contaminants, such as keratins and enzymes used for digestion, as references. Carbamidomethyl cysteine was set as a fixed modification, and protein amino-terminal acetylation and oxidation of methionine were set as variable modifications. The MS/MS tolerance was set to 20 ppm, and three missed cleavages were allowed using trypsin/P as enzyme specificity. Peptide, site, and protein FDR based on a forward-reverse database were set to 0.01. The minimum peptide length was set to 7, and the minimum number of peptides for the identification of proteins was set to one, which must be unique.

Values were median normalized. Missing values in the control (gfp or wildtype) samples were imputed. The Student statistical T-tests (for intensity and iBAQ) were performed after imputation of missing values in the gfp or wild-type samples.

In order to enlarge the number of hits, we also added a less stringent p-value set at a threshold <0.05, which was not adjusted for multiple testing with an FDR<0.05 (as we lost lots of candidates after FDR adjustment for the CoIP-MS).

¨

The ENCODE Project Consortium, Dunham, I., Kundaje, A., Aldred, S.F., Collins, P.J., Davis, C.A., Doyle, F., Epstein, C.B., Frietze, S., Harrow, J., Kaul, R., Khatun, J., Lajoie, B.R., Landt, S.G., Lee, B.-K., Pauli, F., Rosenbloom, K.R., Sabo, P., Safi, A., Sanyal, A., Shoresh, N., Simon, J.M., Song, L., Trinklein, N.D., Altshuler, R.C., Birney, E., Brown, J.B., Cheng, C., Djebali, S., Dong, X., Dunham, I., Ernst, J., Furey, T.S., Gerstein, M., Giardine, B., Greven, M., Hardison, R.C., Harris, R.S., Herrero, J., Hoffman, M.M., Iyer, S., Kellis, M., Khatun, J., Kheradpour, P., Kundaje, A., Lassmann, T., Li, Q., Lin, X., Marinov, G.K., Merkel, A., Mortazavi, A., Parker, S.C.J., Reddy, T.E., Rozowsky, J., Schlesinger, F., Thurman, R.E., Wang, J., Ward, L.D., Whitfield, T.W., Wilder, S.P., Wu, W., Xi, H.S., Yip, K.Y., Zhuang, J., Bernstein, B.E., Birney, E., Dunham, I., Green, E.D., Gunter, C., Snyder, M., Pazin, M.J., Lowdon, R.F., Dillon, L.A.L., Adams, L.B., Kelly, C.J., Zhang, J., Wexler, J.R., Green, E.D., Good, P.J., Feingold, E.A., Bernstein, B.E., Birney, E., Crawford, G.E., Dekker, J., Elnitski, L., Farnham, P.J., Gerstein, M., Giddings, M.C., Gingeras, T.R., Green, E.D., Guigó, R., Hardison, R.C., Hubbard, T.J., Kellis, M., Kent, W.J., Lieb, J.D., Margulies, E.H., Myers, R.M., Snyder, M., Stamatoyannopoulos, J.A., Tenenbaum, S.A., Weng, Z., White, K.P., Wold, B., Khatun, J., Yu, Y., Wrobel, J., Risk, B.A., Gunawardena, H.P., Kuiper, H.C., Maier, C.W., Xie, L., Chen, X., Giddings, M.C., Bernstein, B.E., Epstein, C.B., Shoresh, N., Ernst, J., Kheradpour, P., Mikkelsen, T.S., Gillespie, S., Goren, A., Ram, O., Zhang, X., Wang, L., Issner, R., Coyne, M.J., Durham, T., Ku, M., Truong, T., Ward, L.D., Altshuler, R.C., Eaton, M.L., Kellis, M., Djebali, S., Davis, C.A., Merkel, A., Dobin, A., Lassmann, T., Mortazavi, A., Tanzer, A., Lagarde, J., Lin, W., Schlesinger, F., Xue, C., Marinov, G.K., Khatun, J., Williams, B.A., Zaleski, C., Rozowsky, J., Röder, M., Kokocinski, F., Abdelhamid, R.F., Alioto, T., Antoshechkin, I., Baer, M.T., Batut, P., Bell, I., Bell, K., Chakrabortty, S., Chen, X., Chrast, J., Curado, J., Derrien, T., Drenkow, J., Dumais, E., Dumais, J., Duttagupta, R., Fastuca, M., Fejes-Toth, K., Ferreira, P., Foissac, S., Fullwood, M.J., Gao, H., Gonzalez, D., Gordon, A., Gunawardena, H.P., Howald, C., Jha, S., Johnson, R., Kapranov, P., King, B., Kingswood, C., Li, G., Luo, O.J., Park, E., Preall, J.B., Presaud, K., Ribeca, P., Risk, B.A., Robyr, D., Ruan, X., Sammeth, M., Sandhu, K.S., Schaeffer, L., See, L.-H., Shahab, A., Skancke, J., Suzuki, A.M., Takahashi, H., Tilgner, H., Trout, D., Walters, N., Wang, H., Wrobel, J., Yu, Y., Hayashizaki, Y., Harrow, J., Gerstein, M., Hubbard, T.J., Reymond, A., Antonarakis, S.E., Hannon, G.J., Giddings, M.C., Ruan, Y., Wold, B., Carninci, P., Guigó, R., Gingeras, T.R., Rosenbloom, K.R., Sloan, C.A., Learned, K., Malladi, V.S., Wong, M.C., Barber, G.P., Cline, M.S., Dreszer, T.R., Heitner, S.G., Karolchik, D., Kent, W.J., Kirkup, V.M., Meyer, L.R., Long, J.C., Maddren, M., Raney, B.J., Furey, T.S., Song, L., Grasfeder, L.L., Giresi, P.G., Lee, B.-K., Battenhouse, A., Sheffield, N.C., Simon, J.M., Showers, K.A., Safi, A., London, D., Bhinge, A.A., Shestak, C., Schaner, M.R., Ki Kim, S., Zhang, Z.Z., Mieczkowski, P.A., Mieczkowska, J.O., Liu, Z., McDaniell, R.M., Ni, Y., Rashid, N.U., Kim, M.J., Adar, S., Zhang, Z., Wang, T., Winter, D., Keefe, D., Birney, E., Iyer, V.R., Lieb, J.D., Crawford, G.E., Li, G., Sandhu, K.S., Zheng, M., Wang, P., Luo, O.J., Shahab, A., Fullwood, M.J., Ruan, X., Ruan, Y., Myers, R.M., Pauli, F., Williams, B.A., Gertz, J., Marinov, G.K., Reddy, T.E., Vielmetter, J., Partridge, E., Trout, D., Varley, K.E., Gasper, C., The ENCODE Project Consortium, Overall coordination (data analysis coordination), Data production leads (data production), Lead analysts (data analysis), Writing group, NHGRI project management (scientific management), Principal investigators (steering committee), Boise State University and University of North Carolina at Chapel Hill Proteomics groups (data production and analysis), Broad Institute Group (data production and analysis), Cold Spring Harbor, U. of G., Center for Genomic Regulation, Barcelona, RIKEN, Sanger Institute, University of Lausanne, Genome Institute of Singapore group (data production and analysis), Data coordination center at UC Santa Cruz (production data coordination), Duke University, E., University of Texas, Austin, University of North Carolina-Chapel Hill group (data production and analysis), Genome Institute of Singapore group (data production and analysis), HudsonAlpha Institute, C., UC Irvine, Stanford group (data production and analysis), 2012. An integrated encyclopedia of DNA elements in the human genome. Nature 489, 57–74. https://doi.org/10.1038/nature11247

Yu, G., Wang, L.-G., He, Q.-Y., 2015. ChIPseeker: an R/Bioconductor package for ChIP peak annotation, comparison and visualization. Bioinformatics 31, 2382–2383. https://doi.org/10.1093/bioinformatics/btv145

Zaghet, N., Madsen, K., Rossi, F., Perez, D.F., Amendola, P.G., Demharter, S., Pfisterer, U., Khodosevich, K., Pasini, D., Salcini, A.E., 2021. Coordinated maintenance of H3K36/K27 methylation by histone demethylases preserves germ cell identity and immortality. Cell Rep. 37, 110050. https://doi.org/10.1016/j.celrep.2021.110050
